# Inhibition of pro-atherogenic trimethylamine production from choline by human gut bacteria is not determined by varying chlorogenic acid content in highbush blueberries

**DOI:** 10.1101/2025.06.20.660815

**Authors:** Ashley M. McAmis, Michael G. Sweet, Sydney Chadwick-Corbin, Juanita G. Ratliff, Molla Fentie Mengist, Nahla V. Bassil, Pon Velayutham Anandh Babu, Massimo Iorizzo, Andrew P. Neilson

## Abstract

Elevated blood levels of trimethylamine N-oxide (TMAO) are linked to increased risk of atherosclerosis. TMAO is produced when gut bacteria metabolize quaternary amines such as choline to trimethylamine (TMA), which is converted to TMAO in the liver. Chlorogenic acid (CGA), a phenolic abundant in blueberries, inhibits TMA production. Blueberries may be a TMA- (and TMAO)-lowering food. CGA content in blueberries varies significantly. It remains unclear whether variations in CGA levels influence the TMA-lowering activity of different cultivars. We investigated the impact of blueberry CGA content on inhibition of choline-d_9_ conversion to TMA-d_9_ in our upper gastrointestinal and *in vitro* human fecal model. Preliminary experiments indicated near-total inhibition of TMA-d_9_ production when whole blueberries were tested. Blueberry pulp and sugars recapitulated this complete inhibition, whereas blueberry skins and a fiber had more moderate inhibition. We proceeded with skins (to avoid interferences from sugar-rich pulp, which would not be present in the colon *in vivo*) from 20 highbush blueberry genotypes, chosen for extremes in CGA content. CGA in whole berries was 2.6-146 mg/100 g fresh weight, while CGA in skins was 13.6-975 mg/100 g fresh weight. No differences were observed in TMA-d_9_ production among the 4 highest and 4 lowest CGA genotypes in kinetic curves or area under the curve (AUC) values when skin digesta were fermented with choline-d_9_. However, significant differences were observed between all genotypes compared to blank digesta, with ∼19.4.% reduction in TMA-d_9_ AUCs, indicating that skins provides similar TMA-lowering benefits across genotypes. Levels of free CGA in fermenta of skin digesta were 0.05-0.3 μM, >1000-fold lower than the minimum effective dose we observed for pure CGA *in vitro*, suggesting that blueberry CGA content is not a crucial factor for lowering TMA. Fiber also does not account for most of the inhibitory activity of blueberry skins. Studies are needed to confirm this *in vitro* study and understand how blueberries inhibit TMA and potentially TMAO production *in vivo*.

## 1. Introduction

Cardiovascular diseases (CVDs) exert a large impact on populations worldwide, accounting for ∼32% of all deaths ^1^. This multifaceted group of disorders includes conditions such as atherosclerosis, myocardial infarction, and stroke and represents a leading cause of morbidity and mortality ^2^. Atherosclerosis linked diseases are the leading causes of death in the United States ^3^. The prevalence of CVDs necessitates the need for comprehensive research on pharmaceuticals, surgical interventions, and complementary strategies such as dietary and lifestyle changes to mitigate its influence on global health. In recent years, there has been growing evidence that certain gut-microbiota derived metabolites play a role in the development of inflammatory bowel disease, liver cirrhosis, rheumatoid arthritis, and CVDs ^4^. In particular, the metabolite trimethylamine (TMA), produced from choline, L-carnitine, and γ-butryobetaine by gut bacteria containing trimethylamine lyase (TMA-lyase) complexes, serves as a precursor to trimethylamine N-oxide (TMAO). Choline TMA-lyase, which metabolizes choline into TMA, is encoded by the *cutC/D* gene cluster found in specific gut microbiota ^5,6^.

Studies suggest that elevated concentrations of trimethylamine N-oxide (TMAO) in the blood are associated with increased risk of atherosclerosis ^7,8^. Choline is a semi-essential nutrient that the body requires to produce acetylcholine, phospholipids, lipoproteins, etc. and plays a crucial role in membrane and brain development in fetuses ^9^. In addition, choline is metabolized to TMA by gut bacteria. Once released, TMA is absorbed through the intestinal mucosa and travels through the bloodstream via the hepatic portal system, which delivers blood directly to the liver rather than to the heart ^10^. Once it reaches the liver, TMA is oxidized to TMAO by hepatic flavin monooxygenase 3 (FMO3) ^11^.

Currently, there is no FDA-approved drug to control or lower TMA or TMAO levels. The gut microbiota plays a crucial role in TMA metabolism, and the addition of antibiotics decreases the presence of bacteria carrying the *cutC/*D gene, ultimately reducing TMA production ^12^. Although the use of antibiotics effectively reduces TMA production by targeting bacteria with the cut*C/*D gene, it may also disturb beneficial commensal bacteria, potentially leading to adverse health effects ^13,14^. Strategies such as non-lethal TMA lyase inhibition lower TMA and TMAO production and inhibit the onset of atherosclerosis in rodents ^15^. Lowering FMO3 expression or inhibiting its action may provide an effective strategy to decrease TMAO production ^16^. However, TMA exudes an odor described as ‘rotting fish’, some individuals who have a dysfunctional metabolism of TMA may experience trimethylaminuria (TMAU) which causes sweat, breath, urine, or other bodily excretions to smell of fish ^17–19^. The reduction or inhibition of FMO3 may result in a buildup of TMA, potentially leading to TMAU. Reducing dietary intake of choline proves to be an effective strategy to reduce TMA and TMAO levels ^12^. However, this is not a viable strategy, as choline plays a crucial role in the synthesis of phospholipids and neurotransmitters.

There is a growing interest in the TMA- and TMAO-lowering activities of dietary phenolics ^20,21^. Phenolics may reduce TMAO levels by various mechanisms, including reducing the abundance of bacterial genera that convert choline into TMA, or direct inhibition of TMA lyase. Due to the low oral bioavailability of many phenolics, unabsorbed phenolics accumulate in the colon’s lumen, where *cutC/D* (and other TMA lyase) gene-complex containing bacteria are found ^22–24^. Chlorogenic acid (CGA, or 5-caffeoyquinic acid) (**Figure 1**) is a phenolic commonly found in the diet, and our lab previously identified CGA as a potential inhibitor of TMA production ^21^. CGA and CGA-rich foods exhibit anti-inflammatory, antihypertensive, and antioxidant properties that help reduce the risk of cardiovascular disease and improve overall health.

**Figure 1.**
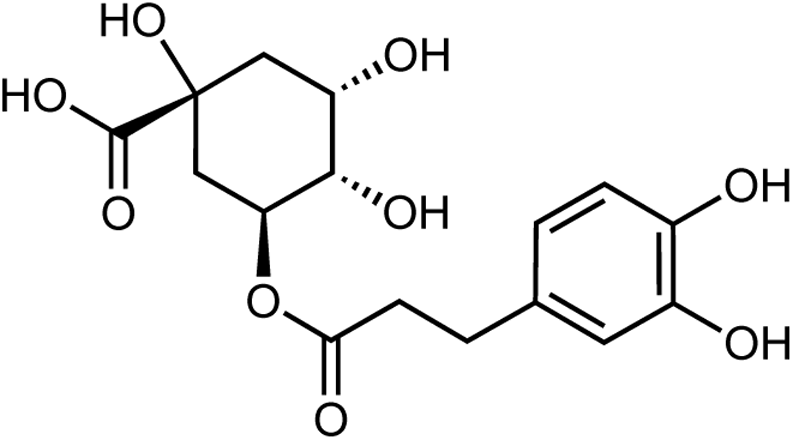
Structure of 5-caffeoyquinic acid (5-*O-*CQA), the most abundant form of chlorogenic acid (CGA) found in highbush blueberries.

CGA stands out as the most predominant phenolic acid in blueberries, accounting for more than 95% of total phenolic acid content in some cases ^25–27^. Although all blueberries contain some level of CGA, the difference between genotypes can vary by >20-fold ^27^. We previously showed that CGA-rich blueberry supplementation reduces TMAO in mice ^28^. However, blueberries contain varying amounts of CGA, so it is challenging to make a general claim that all blueberries offer the same health benefits, as their CGA levels (and other bioactive compounds) may influence these benefits. Additionally, other components may drive the TMA- and TMAO-lowering activities of blueberries.

The overall aim of this study was to evaluate whether CGA is a key bioactive that enables blueberries to inhibit the formation of the gut microbial metabolite TMA (a precursor to the pro-atherogenic compound TMAO) from choline. We hypothesized that CGA content is a primary determinant of the TMA-lowering benefits of blueberries. To test the inhibition properties of CGA and other non-phenolic components of blueberries, an *ex vivo-in vitro* human fecal fermentation model was utilized to evaluate the inhibition of TMA production from choline.

## 2. Methods and Materials

Due to the large number of different methods used in this study, we have provided complete and exhaustive methodological detail for all methods in **Supplementary Information.**

### 2.1 Chemicals

a full list of reagents and materials can be found in **Supplementary Information**. Fecal samples were obtained from OpenBiome (Cambridge, MA, USA). TMA-d_9_ (CAS# 18856-86-5) was obtained from MilliporeSigma (Burlington, MA). Choline-1-^13^C-1,1,2,2, -d_4_, choline-1,2-^13^C_2_, TMA-^13^C_3_-^15^N, and TMA-^13^C_3_-d_9_ were obtained from Cambridge Isotope Laboratories (Tewksbury, MA).

### 2.2 Blueberry Sources

Commercially available frozen (Food Lion, LLC Salisbury, NC 28147) and fresh (SBROCCO International INC. Mount Laurel, NJ 08054) blueberries for method development were obtained from Food Lion, Kannapolis, NC, USA in March 2024, and frozen immediately at −80°C. A 1:1 mixture of these store-bought berries was used for method development, to represent a composite sample of typical blueberries available to consumers for the initial experiments while preserving the limited quantities of blueberry material from the unique genotypes with varying CGA levels for later experiments. Fruits from various blueberry genotypes were obtained from the 2019 and 2024 harvest of a diverse field collection through the United States Department of Agriculture Agricultural Research Service (USDA ARS) National Clonal Germplasm Repository (NCGR), Corvallis OR, USA. Fruit from a bi-parental population (Draper x Jewel, labled as DxJ) were obtained from plants grown by the USDA-ARS Berry Breeding Program at the Corvallis-based Horticultural Crops Production and Genetic Improvement Research Unit (HCPGIRU). Fruits from these two distinct populations were obtained: (1) accessions (labeled as Plant Introduction, PI) belonging to a diversity panel of cultivars/selections where the plants were not directly related and did not share genetic material from the same immediate parental sources, and (2) genotypes belonging to the DxJ bi-parental mapping population where plants shared genetic material from the same parental sources. The accessions and the DxJ genotypes were selected based on chlorogenic acid content evaluated in previous work ^29^ and verified in this study (see results). Also the genotypes from the mapping population were selected based on genotypic data, representing the dominant and recessive allele at a major genetic locus associated with chrologenic acid content ^29^. Blueberries were frozen, shipped overnight on dry ice, and stored at −80°C immediately upon receipt prior to use.

### 2.3 General Gastrointestinal Digestion

Simulated *in vitro* upper gastrointestinal digestions were performed to mimic oral, gastric, and small intestinal digestion following the methodology described by Iglesias-Carres *et al.* ^30,31^. Digestions were scaled from ∼1 serving [150 g fresh weight (FW)] in estimated human upper gastrointestinal volume (2 L) to an equivalent dose for the *in vitro* digestion final volume (15 or 50 mL). “Blank” or “control” digesta was made with saline in place of food material and treated the same as the other treatments. Further details of digestion methods can be found in the **Supplementary Information.**

### 2.4 Anaerobic Fecal Fermentation

#### 2.4.1 Growth Media Preparation

Anaerobic growth media were prepared following the methodology described by Iglesias-Carres *et al.* ^21^. Further information can be found in the **Supplementary Information.**

#### 2.4.2 Fecal Slurry preparation

De-identified fecal samples from healthy donors were obtained through OpenBiome (Cambridge, MA, USA) and stored immediately at −80°C. Fecal slurries were prepared 12 hrs before the start of fermentations (concentrations varied based on experimental parameters). At least two different fecal samples were combined in a ratio of 1 mL fecal sample to 9 mL anaerobic growth media and vortexed until homogenized. The fecal slurry was left uncapped in the anaerobic chamber for at least 12 hrs prior to fermentations. Further details can be found in the **Supplementary Information.**

#### 2.4.3 General Fermentation Conditions

Fermentations took place inside an 855-ACB anaerobic chamber (Plas-Labs, Lansing, MI, USA). The chamber was first purged of oxygen using N_2_ (Airgas, Charlotte, NC, USA) and then mixed gas containing 5% H_2_, 5% CO_2_ and 90% N_2_ (ARC3, Raleigh, NC, USA) until chamber conditions reached H_2_ (2-3%) and O_2_ levels below 20 ppm monitored with a CAM-12 anaerobic monitor (Coy Lab Products, Grass Lake, MI, USA). Once anaerobic conditions were achieved, the heater inside the chamber was turned on to 37°C, and a palladium catalyst (to consume residual O_2_) was placed on top before leaving it to stabilize overnight. Conditions were monitored throughout fermentations. Further information can be found in the **Supplementary Information.**

#### 2.4.4 General Fermentation Procedure

Fermentations were carried out per our established methodology ^21^. We focused exclusively on TMA production from choline using isotopically labeled substrate to eliminate interference by endogenous TMA-lyase substrates. The anaerobic chamber was prepared as described above. Choline-d_9_ was added at a final concentration of 100 μM in all experiments, and choline-d_9_ utilization and TMA-d_9_ production were measured. Digested samples were prepared as described in general gastrointestinal digestions, lyophilized (to remove O_2_) and reconstituted to 1X digesta concentration using filter-sterilized, overnight-sparged PBS 1X inside the chamber. In 1.1 mL 96-well plates, 750 μL of growth media were mixed with 90 μL of choline-d_9_ stock solution (2 mM) in PBS 1X, 600 μL of reconstituted digested samples and 360 μL fecal slurry (1:10 in PBS 1X). Final concentrations were thus 100 μM choline-d_9_, 2% fecal matter, and 33.3 % digesta. Samples containing no digesta were brought to volume with 600 μL of PBS 1X. The start of fermentation (time 0 h) began when fecal slurry was inoculated into the reaction mixture with the substrate. From 0 up to 30 h, a 100 μL sample was collected at various time points, combined with 100 μL of acetonitrile (ACN), and frozen immediately at −80°C. Further details can be found in the **Supplementary Information.**

### 2.8 Phenolic Fraction Extraction

A phenolic extraction was performed to generate a blueberry fraction representing phenolic compounds in an average serving of blueberries (∼150 g) for use in fermentation. Whole blueberries (1:1 mix of locally available store-bought fresh and frozen berries) were homogenized, weighed, lyophilized, weighed and frozen at −80°C. Lyophilized material representing a serving size of fresh weight blueberry was combined with an extraction mixture containing acetone, water, and acetic acid (70:29.5:0.5) and vortexed, blended for 1 min using a polytron (VWR 200), sonicated for 5 min in a water bath at 50 °C, and centrifuged (3248 x g, 10 min). The steps were repeated seven times until the supernatant was clear. The supernatant was collected from the extractions, pooled, and concentrated using a rotary evaporator at 45 °C. The extract was frozen at −80°C and lyophilized. The lyophilized extract was weighed and resuspended in acidified mill-Q water (0.1% formic acid). The resuspended extract material was eluted using Diaion HP-20 resin and a 24/40 column to remove and purify phenolic compounds, then transferred into a round-bottom flask and concentrated using a rotary evaporator. The extract material was frozen at −80°C and lyophilized. Further information can be found in the **Supplementary Information.**

### 2.9 Plant Phenolic Extraction and Analysis

Extraction of blueberry material (skin, pulp and whole fruit) was performed. The Folin assay was carried out to estimate total phenolics in the plant extracts. To quantify the relative amounts of CGA in plant material, a solid phase extraction (SPE) followed by analysis using UPLC-MS/MS was utilized following the methodology described by Mengist *et al*.^32^. Further information can be found in the **Supplementary Information.** CGA MS/MS parameters and a representative UPLC-MS/MS chromatogram are shown in **Supplementary Table 1** and **Supplementary Figure 1**, respectively.

### 2.10 Digesta and Fermenta CGA Analysis

Digesta samples were combined 1:1 with 0.1% formic acid in ACN, vortexed, sonicated for 5 min, then centrifuged (10 min at 17,000 x g). The supernatant was filtered through a 0.2-micron PTFE filter before LC-MS analysis. Fermenta samples were combined 1:1 with ACN after sample collection during fermentation. To acidify these samples, 10 μL of 0.425% formic acid in ACN was mixed with 75 μL of the fermenta sample. Samples were filtered through AcropreAdv 0.2 μm WWPTFE 96-well filtering plates (Pall Corporation, Port Washington, NY, USA) using centrifugation (10 min at 3428 x g) before LC-MS analysis. Further information can be found in **Supplementary Information.**

### 2.11 BacTiter-Glo ATP Assay

Bacterial viability was quantified by luminescence using a commercial BacTiter-Glo Microbial Cell Viability Assay (Promega, USA). The reagent was prepared by mixing equal amounts of the provided BacTiter-Glo substrate and BacTiter-Glo buffer and leaving to rest for at least 15 min at room temperature. A total of 100 μL of the fermenta sample was plated on a white 96-well opaque round-bottom plate and centrifuged (500 x g, 5 min) to remove interfering material. The pellet was resuspended in 100 μL of PBS 1X, combined with 100 μL BacTiter-Glo reagent, and covered for 5 min before being placed in the plate reader (SpectraMax iD3, Molecular Devices, USA). Luminescence was read (Shake: 10s, integration: 2000, read height: 0.50mm) and results recorded. This test is used to determine whether the treatment’s inhibitory effects on choline-d_9_ utilization and TMA-d_9_ production are not a result of low cell viability or general cytotoxicity. Validation of the assay is described in the **Supplementary Information**. BacTiter-Glo validation and performance data are shown in **Supplementary Table 2** and **Supplementary Figures 2-4.**

### 2.12 Extraction and Quantification of Choline-d_9_ and TMA-d_9_

Sample extraction and quantification of choline-d_9_ and TMA-d_9_ using isotope dilution were carried out as described by Iglesias-Carres et al. ^33^. Two internal standards were available for use, with only one utilized per experiment (TMA-^13^C_3_-^15^N or TMA-^13^C_3_-d_9_, and choline-1-^13^C-1,1,2,2, -d_4_ or choline-1,2-^13^C_2_). Further information can be found in the **Supplementary Information.** Structure and reactions schemes for analytes and internal standards are shown in **Supplementary Figure 5.** MS/MS parameters and presesentative UPLC-MS/MS chromatograms for analytes and internal standards are shown in **Supplementary Table 3** and **Supplementary Figures 6-11**, respectively.

### 2.13 Statistical Analyses

Prism 10.4.1 (GraphPad, La Jolla, CA) was used for statistical analyses. One-way ANOVA was used for experiments with one independent variable (treatment effects). Two-way ANOVA was used for experiments with multiple independent variables, as well as choline-d_9_ and TMA-d_9_ kinetic curves (significant main effect/interaction). Mixed effects models were used instead of 2-way ANOVA in the event of missing values. If a significant treatment effect (1-way ANOVA) or main effect/interaction (2-way ANOVA) (P < 0.05) was noted, Tukey post hoc test or Sidak’s multiple comparison test was used to assess the significance of the difference between means (in the case of 2-way ANOVA, or mixed effects models, one family per variable). If kinetic curve calculated values were negative, a zero was imputed for the areas-under-the-curve (AUCs) calculations. Significance was defined *a priori* as α < 0.05.

## 3. Results

### 3.1 Experiment 1: Determination of TMA-d_9_ lowering activities of blueberries with high and low CGA content

Based on previously reported CGA content in the DxJ blueberry bi-parental genetic mapping population (2017-2019 crop years, **Supplementary Figure 12**), eight blueberry genotypes with large differences in CGA content were selected ^27^. Four high CGA genotypes (170-250 mg/100 g FW) containing the dominant allele were compared to four low CGA genotypes (17-32 mg/100 g FW) containing the recessive allele were selected for further study. Simulated digestion was performed at a dose equivalent to 133 g of fresh blueberry consumed by an adult human. Digesta were diluted 3X and fermented for 24 hrs with fecal bacteria and choline-d_9_. Near-complete inhibition of TMA-d_9_ production was observed by all eight genotypes (**Figure 2A**), which we have not previously observed with coffee, tea, cocoa ^30^, artichokes or inulin ^31^. Close examination of TMA-d_9_ appearance kinetics (**Figure 2B**) suggests that differences between CGA genotypes may be present, which might increase if the assay were extended. Components such as sugar or fiber may also have reduced TMA-d_9_ production *in vitro* ^30,34^. Previous work by Bresciani *et al.* showed that sugar, which would not be present in the colon after upper gastrointestinal absorption, can significantly inhibit TMA formation *in vitro* ^34^. This would be experimental interference, rather than a physiologically relevant effect.

**Figure 2.**
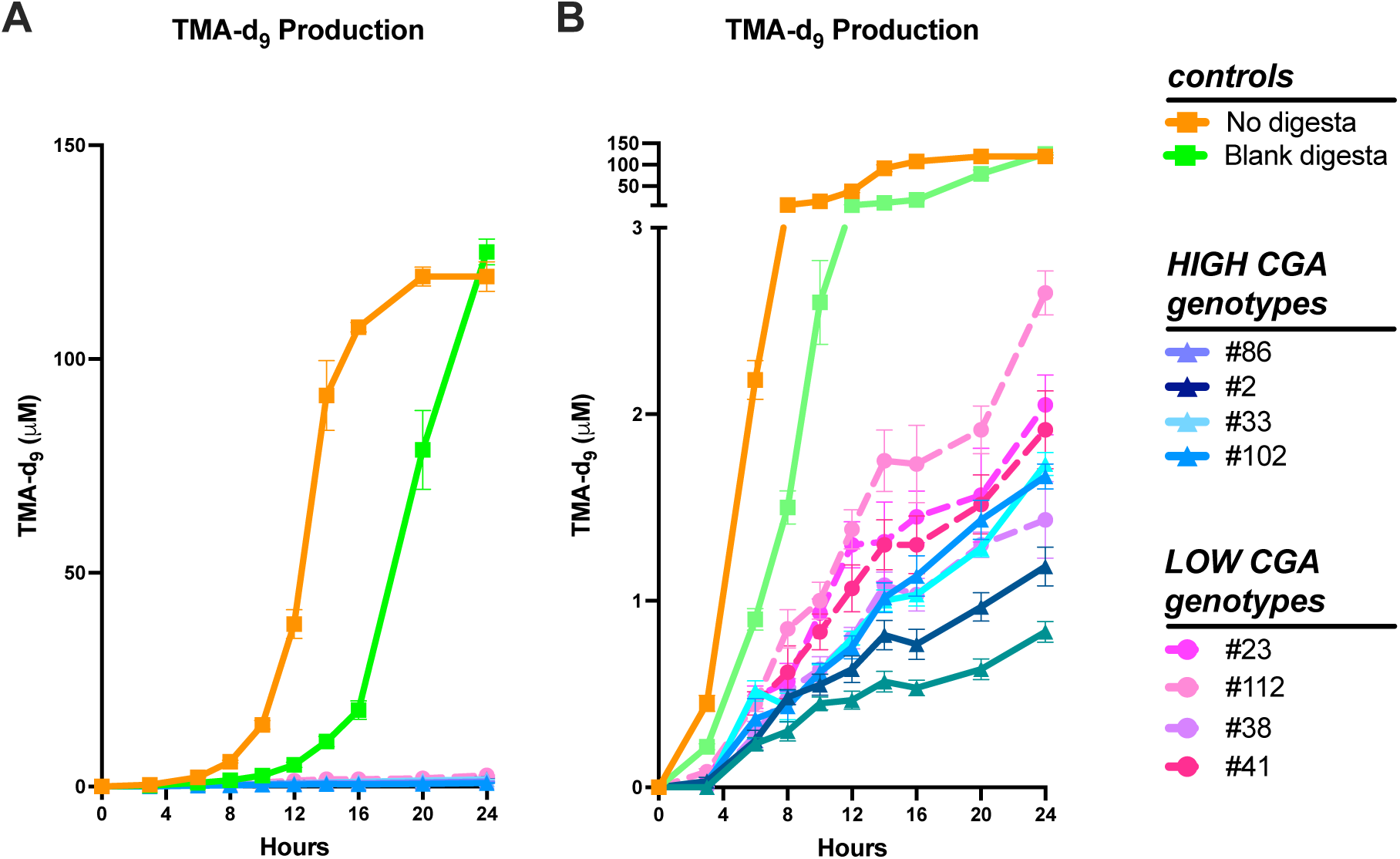
Kinetics of TMA-d_9_ production in fecal fermentations with choline-d_9_: No digesta, Blank digesta, or Digesta from highbush blueberry genotypes from the bi-parental mapping population were added. TMA production displayed without a broken axis (A) and with a broken axis (B). Data represent mean ± SEM from *n*=6 replicates.

### 3.2 Experiment 2: Determination of the impacts of CGA and major fruit components (sugar, fiber, skin, pulp) of blueberries on TMA production

We postulated that sugar, from blueberry pulp, may be responsible for the near-total inhibition of TMA-d_9_ production by blueberries in Expeiment 1. Given the nature of our *in vitro* digestion model, removing sugar is not possible without also eliminating other water-soluble compounds, such as CGA, that are not well-absorbed (and would thus be present in the colon *in vivo*). To identify potential blueberry interferences, whole blueberry, sugar (mixture of glucose, fructose, and sucrose), fiber (mixture of cellulose, hemicellulose and pectin), and phenolic fractions (each matching the amount and composition in the whole berry, **Supplementary Table 4**) were digested and used in an extended fermentation (30 h). Further information can be found in the **Supplementary Information.**

BacTiter-Glo measurement of viable bacteria is presented in **Supplementary Figure 13A**. There is no evidence of broad cytotoxicity, but there were differences between treatments. Kinetics of choline-d_9_ use and TMA-d_9_ production from the fermentations are presented in **Figures 3A-B**, respectively. Whole blueberry digesta and sugar fraction digesta treatments exhibited essentially no choline-d_9_ breakdown or TMA-d_9_ production, as seen in areas-under-the-curve (AUCs) of the choline-d_9_ (**Figure 3C**) and TMA-d_9_ (**Figure 3D**) kinetics. This supports our hypothesis that sugar in the whole blueberry is responsible for the observed near-total TMA-d_9_ inhibition, as sugar (matched in content and composition) mimicked the behavior of whole blueberry. Fiber (which would be present in the colon *in vivo*) did not affect choline-d_9_ use or TMA-d_9_ production, suggesting that fiber does not interfere with the assay.

**Figure 3.**
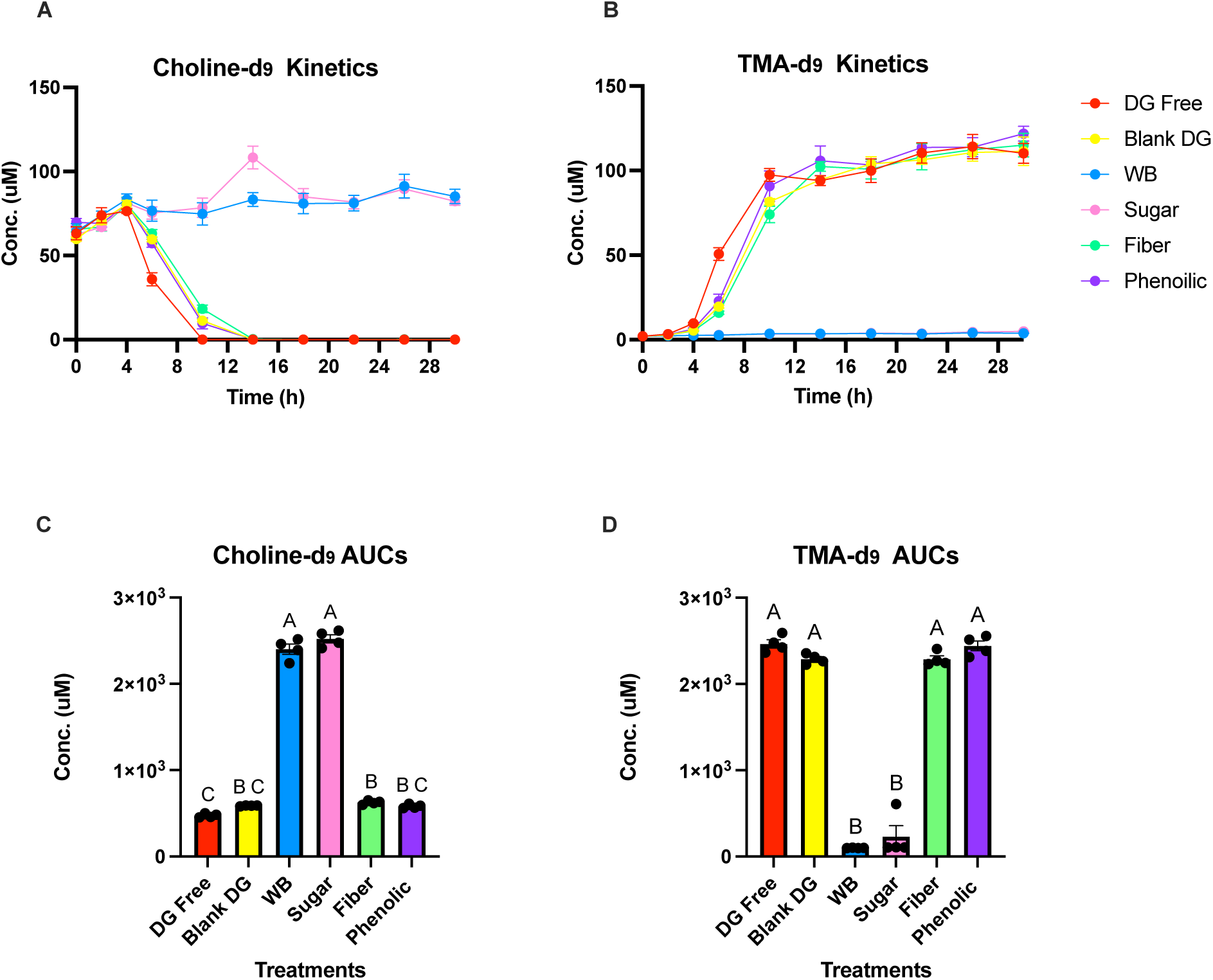
Kinetics of choline-d_9_ use (A) and TMA-d_9_ production (B), and areas under the curve (AUCs) for choline-d_9_ utilization (C) and TMA-d_9_ production (D) in fermentation (0-30 hr) for all treatments including: Digesta free (DG free), Blank DG (blank digesta), Whole blueberry (WB), Sugar, Fiber, and Phenolic fractions. Using one-way ANOVA and Tukey’s multiple comparison to determine statistical differences between treatments. Different letters notate a statistical difference (P<0.05). Data represent mean ± SEM from *n*=6.

### 3.3 Experiment 3: Identification of blueberry components with the greatest TMA inhibition potential

We attempted to further identify blueberry fractions with the most significant TMA-d_9_ lowering effect. We unsuccessfully attempted to remove the sugar post-digestion using sucrose, fructose, and glucose degrading enzymes (data not shown). We elected not to pursue this course further due to the unknown interferences that might arise from sugar degradation products. We then broke the whole blueberry into skin (fiber- and phenolic-rich) vs. pulp (sugar-rich) fractions and compared their inhibitory effects to those of the sugar mixture (glucose, fructose, and sucrose, matching the ratio and content in whole blueberry) and whole blueberry (**Supplementary Table 5**). Total phenolics and CGA content analysis of whole blueberry, skin, and pulp fractions (**Table 1**) confirmed that most phenolics and CGA are located in the skin, as expected. BacTiter-Glo data for blueberry fraction fermentations are presented in **Supplementary Figure 14**. Again, there is no evidence of broad cytotoxicity, but there were differences between treatments. WB and pulp had significantly higher values at both 12 and 24 h. It is interesting that whole blueberry and pulp, but not sugar, increased viable bacterial loads.

**Table 1.** Mean and standard error of the mean (SEM) for commercially available highbush blueberry fractions: Folin Assay (*n*=3) and chlorogenic acid (CGA) content after solid phase extraction (SPE) and LC-MS analysis (*n*=2)

|  | **Folin Assay** |  | **SPE and LC-MS Analysis** |  |
| --- | --- | --- | --- | --- |
| **Material** | **mg GAE/ 100 g whole blueberry<sup>1</sup>** | **SEM** | **mg CGA/ 100 g whole blueberry<sup>1</sup>** | **SEM** |
| Peel | 122.76 | 2.60 | 11.86 | 0.04 |
| Pulp | 40.77 | 0.17 | 6.32 | 0.50 |
| Whole Blueberry | 180.05 | 1.29 | 8.07 | 1.23 |

<sup>1</sup>Whole blueberry values were measured for the bottom row and calculated for the top two rows based on ratios of peel and pulp in the whole blueberry

Kinetics of choline-d_9_ use and TMA-d_9_ production are presented in **Figure 4A-B** for fermentations. Whole blueberry, pulp, and sugar demonstrated near-total inhibition of TMA-d_9_ production. Digesta free, blank digesta, fiber, and skin treatments were significantly different from whole blueberry, pulp, and sugar treatments, and each other. Digesta-free treatments exhibited the quickest choline-d_9_ utilization. Fiber digesta demonstrated the second quickest choline-d_9_ utilization. Blank digesta was similar to fiber. Skin digesta treatments exhibited low choline-d_9_ utilization, though not to the same extent as the whole blueberry, pulp, and sugar treatments. Since the skin contains most of the fiber, the stronger inhibition observed with the skin digeta compared to the matched fiber digesta suggests that some other skin component contributes to inhibition of TMA-d_9_ production.

**Figure 4.**
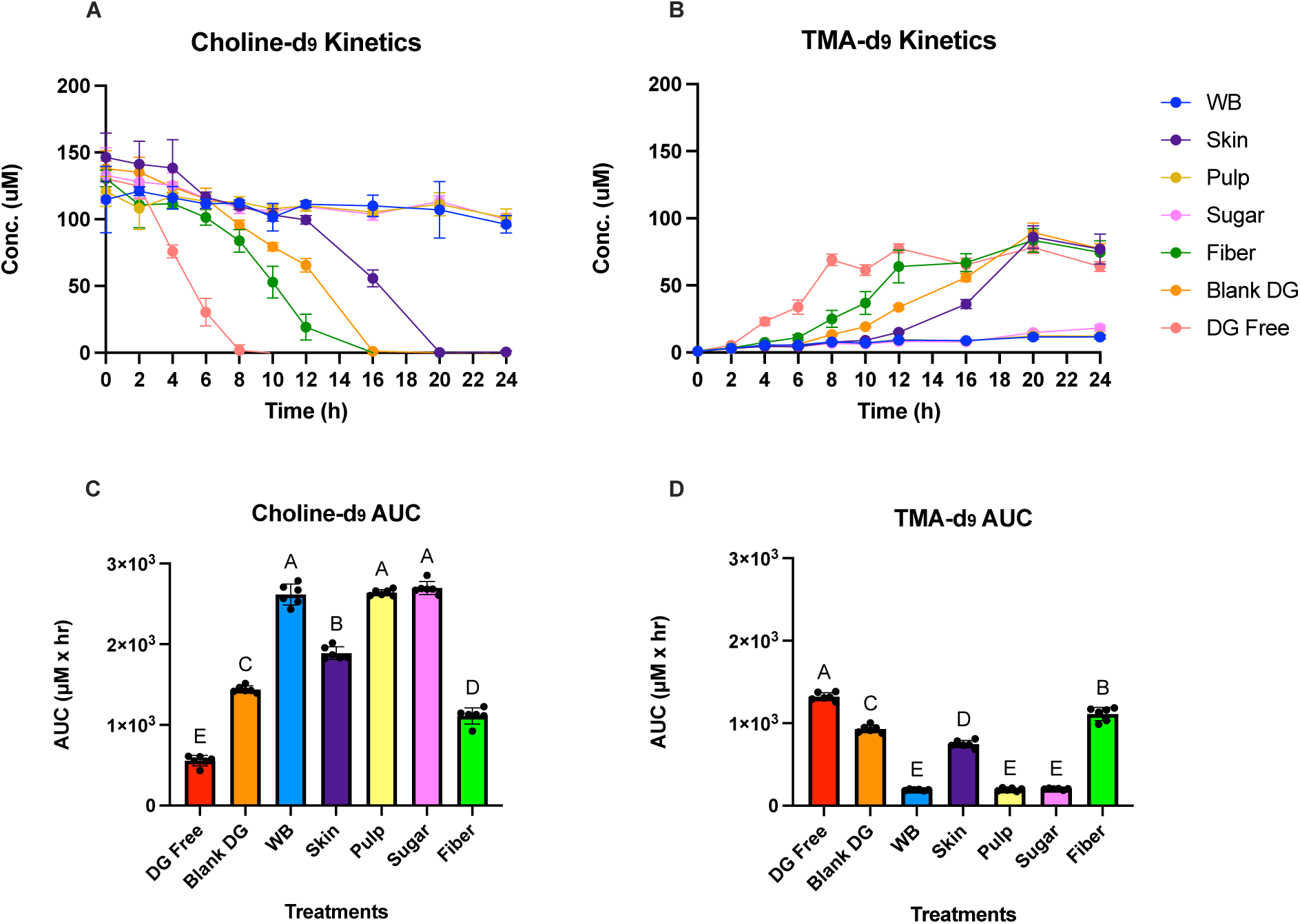
Kinetics of choline-d_9_ use (A) and TMA-d_9_ production (B), and areas under the curve (AUCs) for choline-d_9_ utilization (C) and TMA-d_9_ production (D) in fermentation (0-30 hr) for all treatments of digesta (DG), Whole blueberry (WB), Blank DG, DG free, WB, Peel, Pulp, Sugar, and Fiber. Using one-way ANOVA and Tukey’s multiple comparison to determine statistical differences between treatments. Different letters notate a statistical difference (P<0.05). Data represent mean ± SEM from *n*=6.

AUCs of choline-d_9_ and TMA-d_9_ are presented in **Figure 4C** and **4D**, respectively. Whole blueberry, pulp, and sugar digesta were not statistically different from each other for choline-d_9_ or TMA-d_9_. In contrast, digesta-free, blank, skin, and fiber digesta displayed statistically significant differences in choline-d_9_ and TMA-d_9_ among themselves and were significantly lower compared to whole blueberry, pulp, and sugar digesta. Although the skin contains most of the fiber, distinct AUCs for choline-d_9_ and TMA-d_9_ for skin vs. fiber digesta suggest that a non-fiber component in skin contributes to the observed inhibition. This experiment shows that the extreme inhibitory effects of whole blueberries are mimicked by sugar-rich pulp and matched sugar mixture, strongly suggesting a physiologically irrelevant interference due to sugar, which would not be present in the colon *in vivo*. Therefore, we proceeded with skin-only digestions and fermentations (at doses equivalent to the amount of skin obtained from 1 serving of whole blueberries) to focus on indigestible, non-absorbed materials primarily in the skin.

### 3.4 Experiment 4: Determining whether CGA content correlates with reduced TMA production in multiple blueberry accessions from a genetic diversity population with a large spread of CGA in our ex vivo-in vitro human fecal fermentation model

Whole blueberry concentrations of CGA from 20 blueberry genotypes (selected from 2017-19 data suggesting theywere particularly high or low in CGA content) from the 2024 crop year fruit from two different populations (genetic diversity panel and DxJ bi-parental mapping population) are presented in **Figure 5** and **Table 2**. We selected eight genotypes for fermentation based on the mean CGA content of the whole blueberries (four highest and four lowest genotypes). As skins were used for fermentation based on Experiments 1-3, the eight selected genotypes were analyzed for skin CGA content. For whole fruit (**Figure 6A**), CGA contents in the four highest genotypes (only PI 296339 was different from DxJ 002) were significantly greater than the four lowest genotypes, which were not different from each other. Differences were also observed in skin CGA content (**Figure 6B**), where again the four genotypes with the highest whole fruit CGA had significantly greater skin CGA than the four genotypes with the lowest whole fruit CGA. Interestingly, PI 296399 (the cultivar with the highest whole fruit CGA content), had 3-4x greater skin CGA content than any of the other high CGA genotypes, which were similar. The four lowest genotypes were not different from each other. As the whole blueberry CGA content is thought to be mostly due to CGA in skin, we plotted correlations of whole fruit CGA as a function of skin CGA (**Figure 6C**). While a correlation was observed (r² = 0.492), it was skewed by PI 296399, and the slope approached but did not reach significance (p = 0.053). Without PI 296399 (**Supplementary Figure 15**), the correlation was strong (r^2^=0.945, slope p=0.0002). Another factor determining CGA content is average fruit size. Smaller fruits have more surface area (and thus more skin) per mass unit of whole fruit. We plotted the CGA content as a function of mean fruit size for all 20 genotypes from the 2024 crop year (**Figure 6D**). While r^2^ was moderate (0.2175), the correlation was significant (slope p=0.0382). It warrants noting that fruits of all sizes (∼5-20 mm mean diameter) were high CGA and low CGA genotypes. Interestingly, the distribution of fruit diameters of medium CGA genotypes was tight (∼12-15 mm). These data suggest that skin CGA content and fruit size determine whole fruit CGA content.

**Figure 5.**
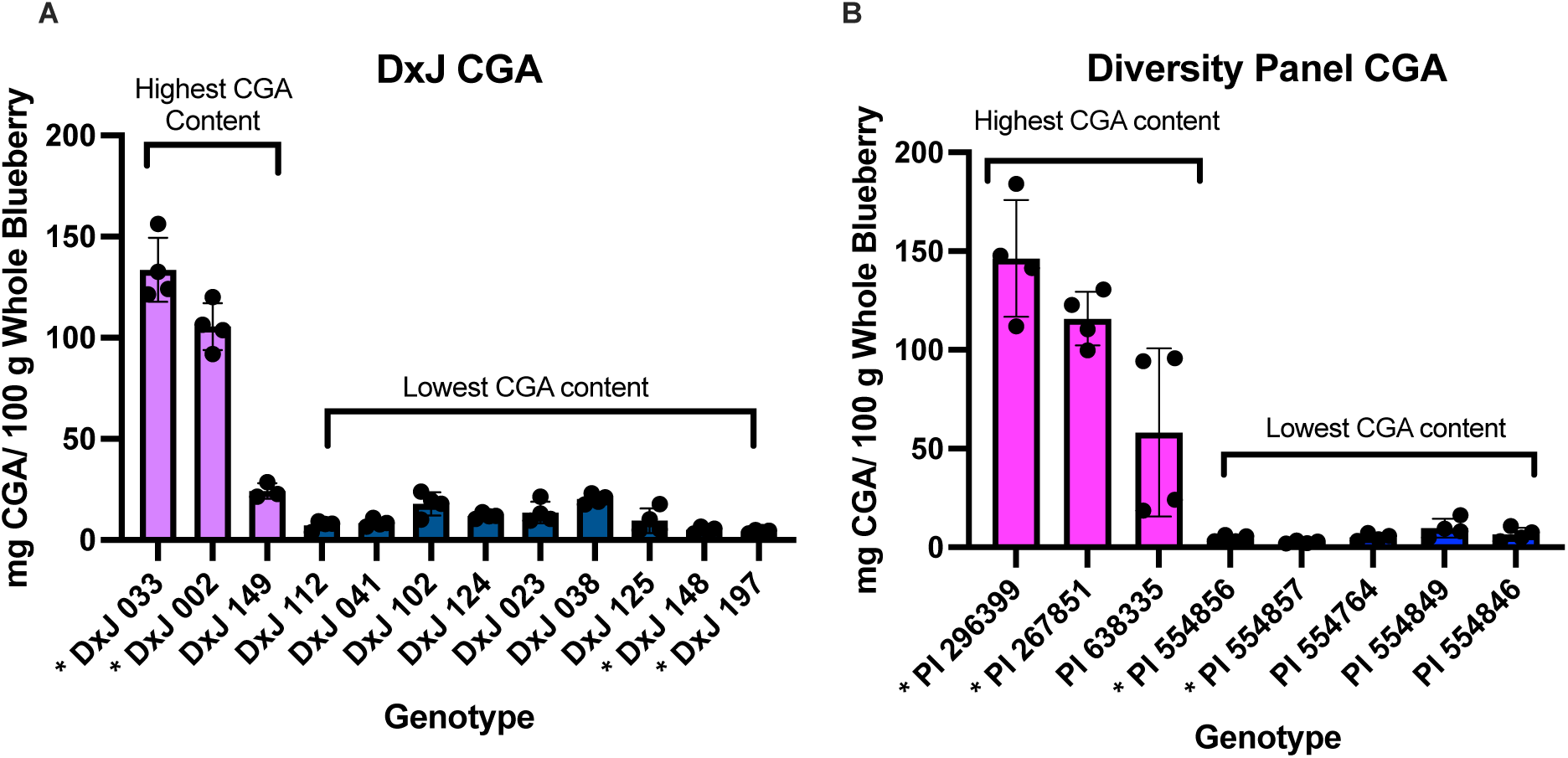
Chlorogenic acid (CGA) content of multiple genetically diverse highbush blueberry genotypes from the summer of 2024. Two (DxJ 197, DxJ 148) of the lowest CGA and two (DxJ 033, DxJ 002) of the highest CGA content blueberries from the biparental mapping population (A) and two (PI554857, PI 554856) of the lowest CGA and two (PI296399, PI 267851) of the highest CGA content blueberries from the diversity panel (B) were selected and noted with an asterisk. Data represent mean ± SEM from *n*=4.

**Figure 6.**
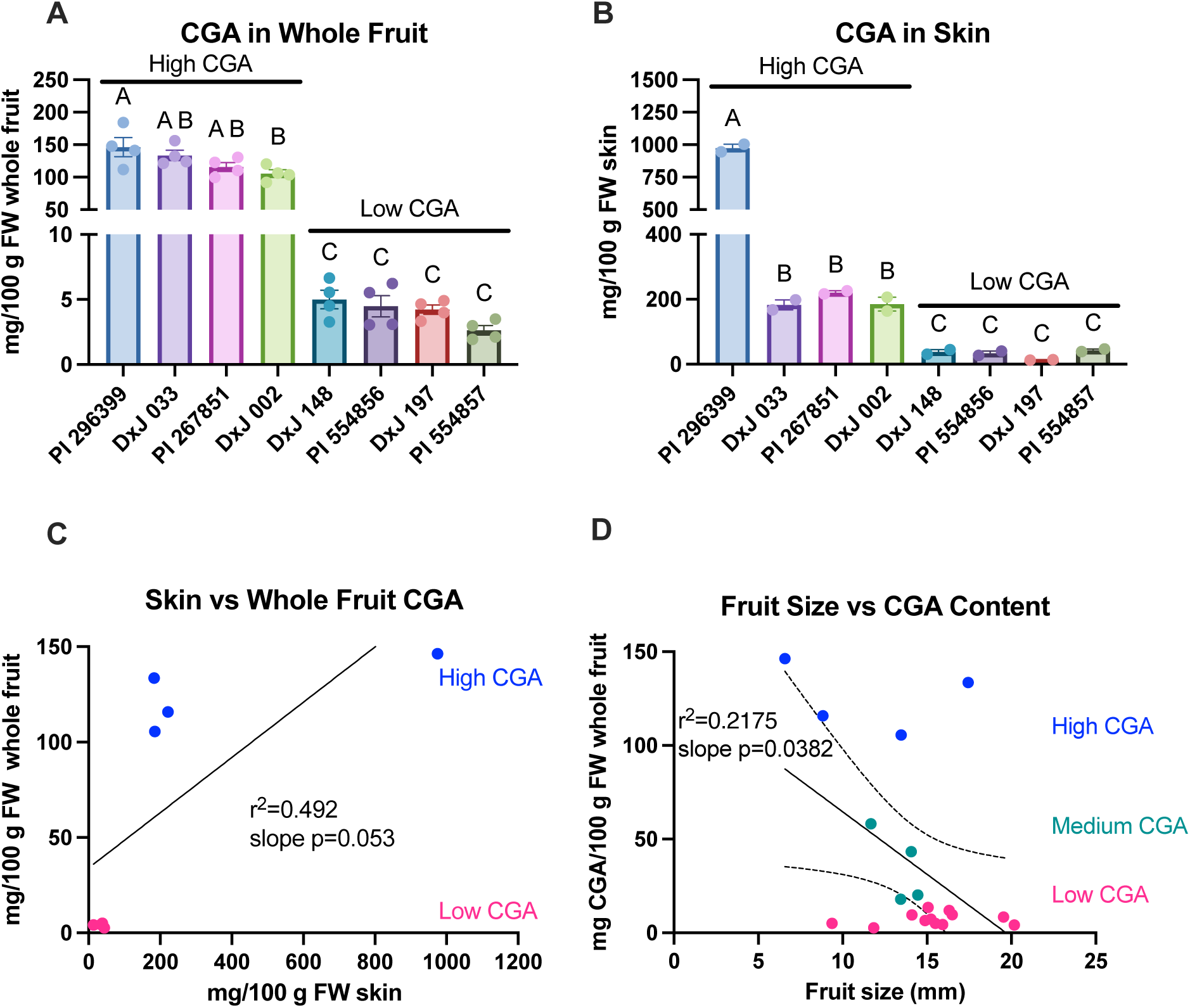
Chlorogenic acid (CGA) content in whole fruit (A) and skins (B) of highbush blueberry genotypes. Correlation of CGA content in skins and whole fruits in the 8 selected genotypes with extreme CGA content (C). Correlations of CGA content with fruit size in all genotypes. Data represents mean ± SEM for *n*=4 (A) or *n*=2 (B). Dots represent means (C and D). FW: fresh weight.

**Table 2.** Average chlorogenic acid (CGA) content in 20 highbush blueberry genotypes (*n*=4)

| **Genotype<sup>1</sup>** | **mg CGA/ 100 g whole blueberry** | **SEM** |
| --- | --- | --- |
| DxJ 197 | 4.22 | 0.36 |
| DxJ 148 | 5.00 | 0.72 |
| DxJ 112 | 7.28 | 1.17 |
| DxJ 041 | 8.39 | 0.90 |
| DxJ 125 | 9.50 | 3.01 |
| DxJ 124 | 11.95 | 0.68 |
| DxJ 023 | 13.57 | 2.69 |
| DxJ 102 | 17.88 | 2.88 |
| DxJ 038 | 20.17 | 1.22 |
| DxJ 149 | 43.35 | 19.16 |
| DxJ 002 | 105.56 | 5.79 |
| DxJ 033 | 133.52 | 7.92 |
| PI 554857 | 2.63 | 0.36 |
| PI 554856 | 4.47 | 0.82 |
| PI 554764 | 5.07 | 0.89 |
| PI 554846 | 6.56 | 1.69 |
| PI 554849 | 9.67 | 2.37 |
| PI 638335 | 58.16 | 21.31 |
| PI 267851 | 115.83 | 6.79 |
| PI 296399 | 146.29 | 14.82 |

<sup>1</sup>DxJ: Biparental mapping population, PI: genetic diversity panel population

Digestions and fermentations of these eight genotypes were carried out (**Supplementary Table 6**). BacTiter-Glo data are presented in **Supplementary Figure 16**. There is no evidence suggesting that any treatment caused a cytotoxic effect that would lead to the perceived effect of TMA-d_9_ inhibition. All genotypes had lower viability values than the controls (but no differences between each other) at 12 h. All genotypes had lower viability than digesta-free, but all were similar to or higher than blank digesta, at 24 h.

Kinetics of choline-d_9_ use and TMA-d_9_ production are presented in **Figure 7A-B**, respectively. Digesta-free treatments exhibited the quickest choline-d_9_ utilization, due to the lack of digesta or skin material to inhibit the reaction. Blank digesta demonstrated the second quickest choline-d_9_ utilization. Enexpectedly, choline-d_9_ utilization was similar across all eight genotypes, regardless of CGA content, with a significant decrease between 8 and 12 hr. While no differences were observed among skins, their inhibition was higher than the controls.

**Figure 7.**
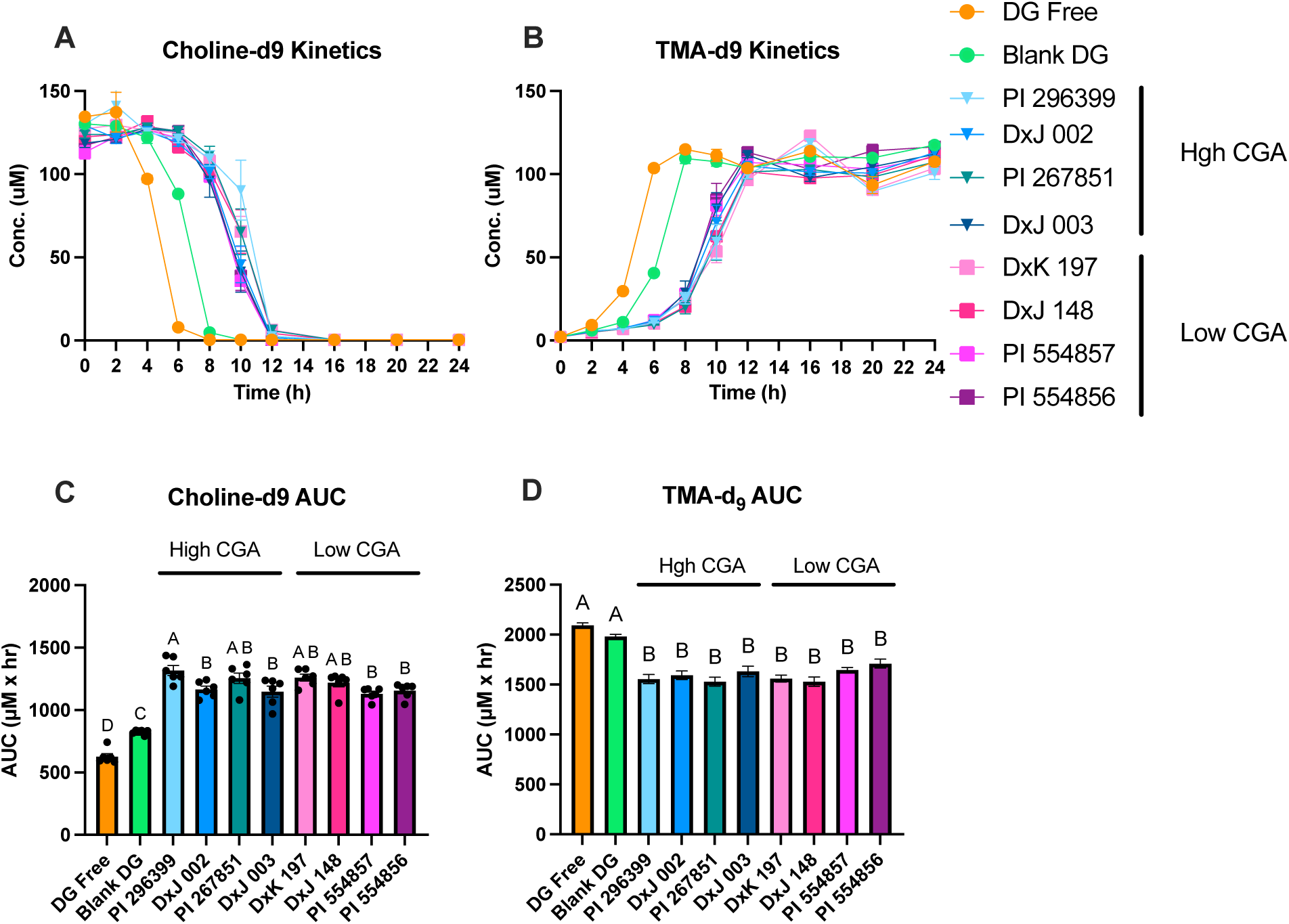
Kinetic for choline-d_9_ utilization (A) and TMA-d_9_ production (B) and area-under-the-curve (AUC) values for choline-d_9_ utilization (C) and TMA-d_9_ production (D) during *in vitro* fecal fermentation of choline-d_9_ with *in vitro* digesta from highbush blueberry skins with varying chlorogenic acid (CGA) levels (0-24 hr). Treatments were compared using one-way ANOVA and Tukey’s multiple comparison. Values not sharing a common superscript letter are significantly different (P<0.05). Data represent mean ± SEM from *n*=6.

AUCs of choline-d_9_ and TMA-d_9_ are presented in **Figure 7C-D**, respectively. Choline-d_9_ AUCs again indicate that skins exhibited a greater inhibitory effect than controls. While there are some differences in choline-d_9_ AUCs between genotypes, these are small and do not suggest a clear relationship to CGA content. TMA-d_9_ AUCs for all genotypes are all lower than controls but not different from one another, suggesting that CGA content does not play a significant role but that all skins exhibited inhibition potential greater than the digestive background.

One confounding variable in our design is that we selected genotypes based on whole fruit CGA content (**Figures 5** and **6A**), but we actually used skins from those genotypes (**Figure 6B**) as opposed to whole fruit in the digestion and fermentation, for the reasons explained above. We employed the same mass of skin from each cultivar (∼20 g skin from a 150 g FW serving of whole fruit, based on experiments to determine the ratio of skin and pulp in whole berries, **Supplementary Table 7**) despite differences in average fruit sizes between genotypes (and thus different skin/pulp ratios, which would result in different amounts of skin between genotypes per 150 g serving of whole berries). This could have unintentionally affected the amount of CGA in each digestion and fermentation. To further assess the potential role of blueberry CGA in reducing bacterial TMA production, we performed linear regression analysis of choline-d_9_ and TMA-d_9_ AUCs (representing total choline-d_9_ and TMA-d_9_ utilization and production, respectively_)_ as a function of both whole fruit CGA content (**Supplementary Figure 17A-B**) and skin CGA content (**Supplementary Figure 17C-D**). Correlations of choline-d_9_ and TMA-d_9_ AUCs with whole fruit CGA are based on the premise that the majority of the CGA in the whole fruit comes from the skin, but ignore the reality that we employed skin, not whole fruit, and that various blueberry genotypes have distinct sizes and thus surface (skin) areas per 150 g serving. Such correlations (**Supplementary Figure 17A-B**) were extremely poor, with both r^2^ < 0.1. Correlations of choline-d_9_ and TMA-d_9_ AUCs with skin CGA more accurately reflect the experimental conditions, as equal masses of skin were used. Such correlations (**Supplementary Figure 17C-D**) were stronger than for whole fruit CGA, but neither had r^2^ >0.25. It should be noted that the distributions of skin CGA concentrations were different from those of whole fruit (**Figure 6A-B**). However, our skin analysis from the eight genotypes selected for fermentations confirms that there were major differences in skin CGA content (**Figure 6B**), albeit not as different as amounts in whole fruit (**Figure 6A**). While whole fruit CGA content was somewhat correlated to skin CGA content (r^2^=0.492, **Figure 6C**), the relationship is not linear. This is due to variations in both fruit size and skin CGA content (**Figure 6B, D**). PI 296399 had the highest CGA content of all genotypes in both whole fruit and skin (**Figure 6A, B**), but its skin CGA content was ∼5x greater than that of any other genotypes. In summary, these correlations show that CGA content was not a good predictor of *in vitro* choline-d_9_ utilization or TMA-d_9_ production. Given that the AUCs and kinetics were similar for all treatments (**Figure 7**), it is unlikely to be a compositional variable that strongly predicts differences in activity.

### 3.5 Experiment 5: Analysis of digesta and fermenta CGA content

As our data do not support the hypothesis that differences in CGA result in differences in inhibition of TMA-d_9_ production, we next attempted to identify factors that might have resulted in a lack of difference in CGA available to interact with the bacteria in the fermentations. Bioaccessibility (digestive release from bulk blueberry) and stability factors could have altered the amount of CGA free to interact with TMA-producing bacteria so that differences in apparent CGA concentrations in digesta and fermenta were not significantly different between treatments. Analyses of digesta and fermenta were performed to determine the levels of free CGA. The relative CGA levels in digesta generally mirrored those in whole fruit and skin, with high CGA genotypes containing more free CGA in the digesta (**Figure 8A**). However, bioaccessibility data (percent of free CGA from the skin released into the digesta) revealed that genotypes with lower skin CGA content had higher % release (but still lower absolute levels) of free CGA compared to those with higher skin CGA content (**Figure 8B**). CGA in the fermenta overall reflected that of the genotypes, with those having higher CGA content also showing higher levels of CGA in fermenta, compared to genotypes with lower CGA content (**Figure 9**). However, there was significant variation in free CGA among the four high CGA genotypes, which did not reflect their skin CGA contents (**Figure 6B**), although PI 296399 was the highest in both. There were no differences in free CGA among the four low CGA genotypes, and all were lower than the four high CGA genotypes. The % release, or bioaccessibility, of the genotypes showed that significant CGA is present in bound forms or is not easily extractable, and a greater % release is generally obtained for low CGA genotypes as opposed to high CGA genotypes. Fermenta samples from high CGA genotypes exhibited higher CGA levels than fermenta of low CGA genotypes (**Figure 9**), although the magnitude of differences was smaller than in skins (**Figure 6B**). All fermenta samples exhibited a decline in CGA levels to essentially the same levels as fermentation progressed. Levels of free CGA detected in fermenta of skin digesta were 0.05-0.3 μM, >1000-fold lower than the minimum effective dose we observed for pure CGA *in vitro* (∼2 mM) ^21^.

**Figure 8.**
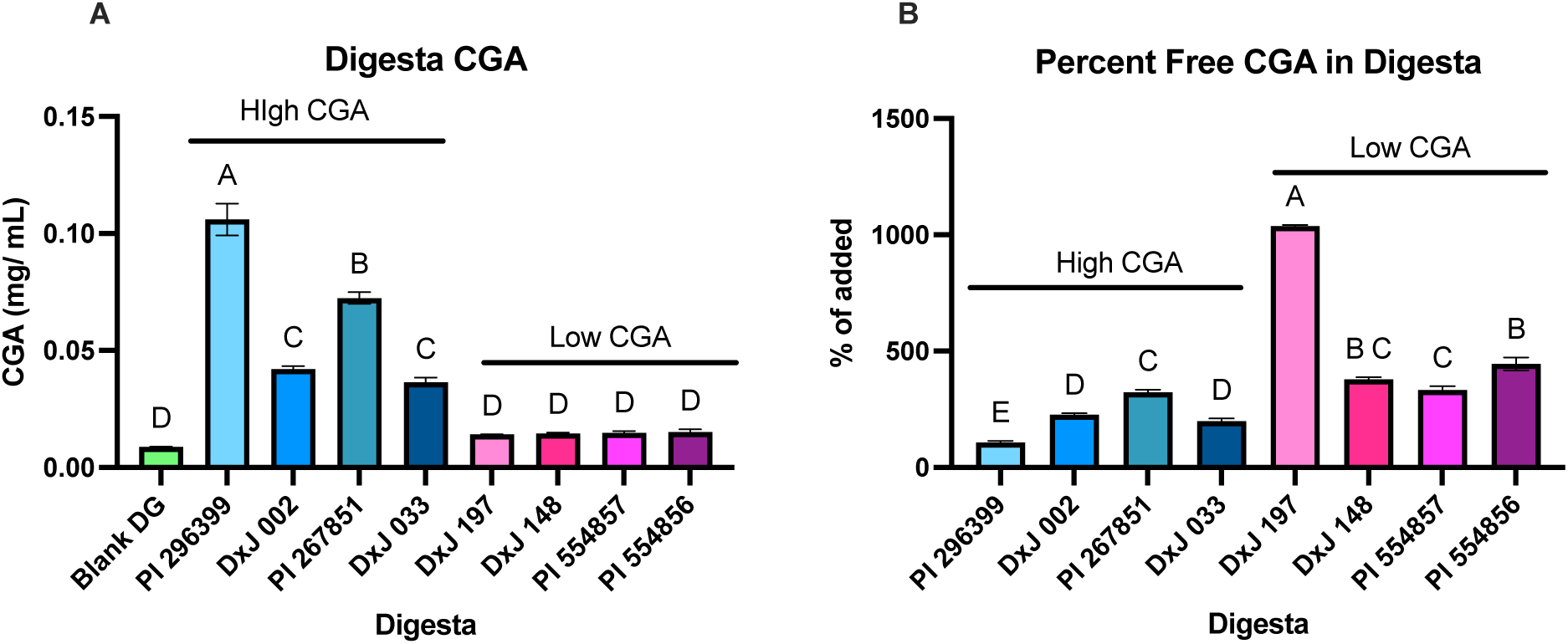
Concentration of free chlorogenic acid (CGA) in whole *in vitro* digesta of highbush blueberry skins (A) and percent CGA released (i.e. bioaccessibility) during digestion (B). Data represent mean ± SEM from *n*=2.

**Figure 9.**
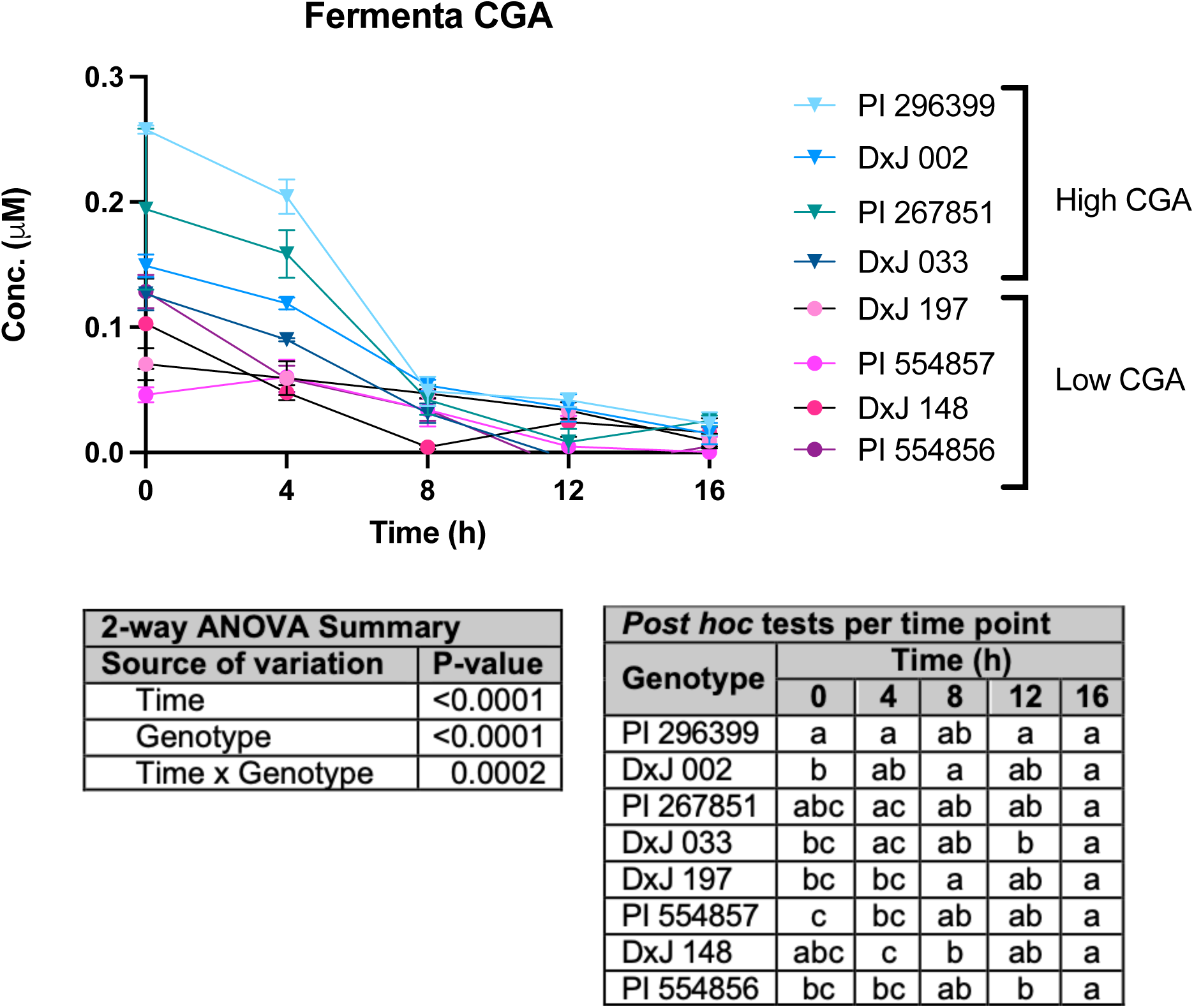
Concentrations of free chlorogenic acid (CGA) in *in vitro* fecal fermenta from highbush blueberry skin digesta from 0-16 h. Data represent mean ± SEM from *n*=3. Tables show the significance of main effects and their interaction in a 2-2qy mixed effects model analysis, and differences among genotypes at each time point (genotypes not sharing a common superscript within the same time point are significantly different by Tukey’s test, P<0.05).

## 4. Discussion

Given the interest in plant bioactives for disease prevention and the mixtures of bioactive phytochemicals in most plants, there is a need to understand which bioactives are essential for the observed activities of a given plant food. One approach to link bioactives with activities is to compare distinct genotypes of the same crop with a range of levels of the bioactive. In the present study, we attempted to determine whether blueberries lower production of the pro-atherogenic microbial metabolite TMA *in vitro*, and whether the level of CGA in blueberries determined TMA-lowering activity. We employed sequential *in vitro* digestion and fecal fermentation to assess conversion of choline-d_9_ to TMA-d_9_ by gut bacteria. We previously studied isolated phenolic compounds ^21,35^, or foods rich in phenolics and/or fiber ^30,31^, but not sugar. Optimizing fermentation parameters for high-fiber, sugar-rich fruits was crucial for identifying the source of the inhibition in blueberries. This was particularly important since Experiment 1 (Figure 2) highlighted the significant inhibitory effects of these highbush blueberry genotypes, surpassing that of any food previously tested in our lab. In Experiment 2, we demonstrated that sugar mimics near-total inhibition of TMA-d_9_ production seen in whole blueberry, but fiber and phenolic extract did not. This suggests, as others have shown ^34^, that sugar interferes with the assay and that the apparent total inhibition by whole blueberry (an initially promising finding) is not physiologically relevant to *in vivo* conditions. Additionally, this showed that major fiber components provide little activity, and perplexingly, neither did a crude phenolic extract. However, these store-bought highbush blueberries had relatively low CGA content compared to the high CGA genotypes used later (**Tables 1-2**) and may have had low levels of other phenolics. The lack of efficacy of the phenolic fraction in these berries may not be indicative of a similar lack of phenolic activity in phenolic-rich berries. In Experiment 3, we compared the activities of skin and pulp to whole blueberry, sugar, and fiber. While skin exhibited less inhibition compared to the whole blueberry, pulp, and sugar treatments, it still showed greater inhibition than the fiber and control treatments. Since the skin contains most of the fiber, the greater inhibition observed with the skin compared to the fiber suggests another skin component is contributing to inhibition of TMA-d_9_ production. We then postulated that, as skin contains most of the phenolics, including CGA, as well as most of the components that will reach the colon *in vivo*, comparing skin of genetically diverse highbush blueberry genotypes with varying CGA content could reveal whether CGA differences result in activity differences. In Experiment 4, we compared skins from high- and low-CGA blueberries. Inhibition potential among all skins was similar, and greater than the controls, but not different between each other.

The average reduction in TMA-d_9_ AUC was 19.4% for the eight genotypes compared to blank digesta. While TMA-d_9_ levels from all genotypes eventually reached the same levels as the controls, this was delayed by ∼ 4 h. *In vivo*, this delay is critical, allowing choline to be absorbed, metabolized to other metabolites, or pass out of the colon. Any of these alternatives will reduce the amount of TMA absorbed and the resulting TMAO concentrations in circulation. Our data suggest that CGA is not the primary factor driving the inhibition in blueberries. Free CGA in the fermenta from physiologically relevant doses of blueberry skins (**Figure 9**) was 1,000-fold more dilute than the minimum effective concentration of pure CGA we previously observed for inhibiting TMA production. Experiment 5 sought to explain the lack of observed differences based on concentrations of free CGA in digesta and fermenta, as the lack of differences in TMA inhibition across genotypes may be due to the lack of variation in CGA concentration once it reached the fermentation. Differences in CGA content in skins were reduced during digestion and further reduced in fermentation. While higher CGA genotypes had higher concentrations in digesta and fermenta, all concentrations were nearly the same by 8 hrs of fermentation.

Our data suggest that blueberry skin significantly reduces generation of TMA by gut bacteria, but that CGA does not drive TMA-lowering activities of blueberries. The compounds responsible for this activity remain unknown. Our data suggest that this inhibition is not due to major blueberry skin fiber components (pectin, cellulose, and hemicellulose), as fiber did not inhibit (but rather promoted) TMA production compared to controls. Thus, the responsible compounds may be other minor fiber compounds or phenolics that we have not yet evaluated. The likely candidates are anthocyanins. We have not yet evaluated the activities of anthocyanins, which are abundant in blueberry skins. Instead of focusing on identifying compositional differences and then testing activity differences based on these compositional differences one by one, the best approach may be to gather a large number (∼50-100) berry samples from many different sources, profile their TMA-lowering activities, then perform broad phenolic profiling on the 10-20% most vs. 10-20% least effective samples.

There are various mechanistic explanations for the observed inhibition of TMA production. Broadly reducing the viability of gut bacteria in a non-specific fashion would lower TMA production ^36^. We employed an ATP-based bacterial activity assay to account for this. The data do not suggest that any component of the blueberry caused cell death leading to TMA-d_9_ inhibition. Furthermore, this is not a likely mechanism as blueberries are widely consumed and are not associated with broad antibacterial activities or adverse gut effects. The observed TMA-lowering activities are likely due to either direct or indirect effects on the activity of *cutC/D*. Potential direct effects include inhibition of choline TMA lyase or reducing its expression. Indirect effects include altering the composition of the gut microbiome by reducing the viability or abundance of *cutC/D*-carrying bacteria. We have previously shown that long-term blueberry supplementation alters the abundance of gut microbiota that are associated with circulating TMAO in experimental mice ^36^. However, given the acute nature of the present study in which the fecal samples are only exposed to blueberry skins for 24 h (and most inhibition is observed within 12 h), we believe that direct effects (reduced *cutC/D* expression and/or inhibition of choline TMA lyase) are the most likely explanation. Both direct and indirect effects may be observed *in vivo*. Future experiments must identify the mechanism of action. Reduced *cutC/D* expression could be investigated with reverse-transcriptase (RT)-qPCR to detect *cutC/D* mRNA or alternatively using Western blotting to detect choline TMA lyase protein. These studies will likely require more enriched cultures of known *cutC/D* carriers, such as *Proteus mirabilis*, as opposed to dilute *cutC/D* in fecal samples ^15^. Studying the inhibition of choline TMA lyase requires the elimination of all other potential mechanisms by excluding viable cells. This could be done with recombinantly expressed enzyme or using a lysed, non-viable *cutC/D* carrier ^15^.

The present study has strengths that enhance its validity. First, the *in vitro* digestion model incorporates physiologically relevant concentrations of digestive enzymes and conditions designed to simulate upper gut digestion. The use of physiologically relevant concentrations of test material maximizes translation portential. Our *in vitro* digestions used 1 serving of whole berries (or equivalent amounts of phenolic extract, fiber, sugar, skin or pulp from 1 serving) per 2000 mL total upper gastrointestinal volume and scaled these ratios to our *in vitro* final volume. When transferring digesta to fermentations, we employed digesta at 33% of the fermentation volume. This is a conservative dilution, which still resulted in significant reductions in TMA-d_9_ production. However, actual concentrations of inhibitory compounds in the colon may be higher, as a critical function of the colon is to extract water from the digesta, thereby concentrating it ^37^. The addition of digesta, rather than whole food, into the fermentation is more representative of a real human system, as it accounts for the presence of digestive enzymes and related metabolites. However, as discussed above, one limitation of this approach for macronutrient-rich foods is that sugars, amino acids and fatty acids that would be absorbed *in vivo* in the stomach or small intestine prior to colonic fermentation are not removed in this model. The *in vitro* colon-only model allows for isolation of TMA production from other mechanisms that would affect assessment of bioactivity, such as absorption of choline and TMA, and gut transit. The use of isotopically labeled choline-d_9_ substrate and measurement of TMA-d_9_ reduces interference from background substrates and products ^21^. We employed fecal samples from OpenBiome, a trusted provider of samples with a healthy and functioning microbiome. Since gut bacteria expressing the *cutC*/D gene cluster can vary among individuals, pooling multiple OpenBiome fecal samples ensures a more diverse population of gut bacteria for each TMA assay. The use of “blank” digesta controls (containing digestion reagents and enzymes, but no blueberry) in the fermentation accounts for background inhibition and endogenous choline from the digesta. Additionally, use of an isotopically labeled substrate in fermentation ensures that the metabolites of interest are accurately measured and tracked, considering the potential presence of endogenous choline and TMA from the digesta. We employed sugar and fiber controls designed to closely reflect the sugar and fiber composition of a serving size of blueberries. The sugar mixture included sucrose, glucose, and fructose in ratios typical of those found in blueberries, while the fiber blend contained hemicellulose, cellulose, and pectin in proportions similar to those in a serving of the fruit. Although literature on blueberry fiber composition is limited, the ratios used in these experiments were believed to accurately reflect the fiber content of blueberries and closely replicate the sugar content. Finally, the key strength of this study is the final experiment, in which eight genotypesof berries were assessed. Too often, experiments are done in a single cultivar of unknown genetic background, and the translatability of the results to consumers is unknown. The blueberries used in the final experiment represented a broad range of highbush genotypes, reflecting a variety of phenolic compositions, sugar content, and fiber levels. Any differences observed among these genotypes are likely due to genetic variations, as they were all grown and harvested in the same climate by the USDA ARS NCGR in Corvallis, OR, USA. Growing in the same climate helps eliminate potential differences caused by factors like water availability, predators, or nutrient competition. The lack of variation in inhibition potential across the peels of the four high and four low genotypes is encouraging for highbush blueberry consumers, as it suggests that the benefits are not limited to a specific cultivar, making it easier for people to access the same benefits regardless of the type of blueberry available.

Our findings should be considered in the context of the following limitations. This work was conducted *in vitro*, which poses limitations. The absorption and conversion of TMA to TMAO and TMAO excretion were not evaluated. These processes affect TMAO in circulation. While we believe that inhibition of TMA formation is the key mechanistic target for lowering TMA, *in vivo* studies are needed to account for all factors that control TMAO levels in circulation. Furthermore, we studied production of TMA from choline exclusively (via *cutC/D*) by adding exogenous labeled choline-d9 and monitoring TMA-d_9_ production. Other precursors (carnitine and betaine) are converted to TMA by TMA lyases encoded by the *cntA/B* and *yeaW/X* gene clusters respectively^7,8,38,39^. Therefore, the present results are only applicable to TMA arising from choline via *cutC/D*. The impact of blueberry skins on TMA levels due to other precursors and their respective TMA lyases remains unknown and merits study in the future. Other studies have previously examined the potential inhibition of TMA formation from carnitine by dietary bioactive compounds^34,40,41^, and the differences in TMA production from various TMA lyases and their respective precursors could mean that a compound that is an effective inhibitor for one pathway may not be effective for another.

As mentioned above, another major challenge is addressing the limitations of *in vitro* digestion and fermentation when using whole fruits, as sugar is not absorbed in the model prior to fermentation. As we and others have demonstrated ^34^, sugar almost completely inhibits microbial TMA-d_9_ generation, but this has little physiological relevance, as sugar would not be present in the colon after upper gastrointestinal digestion. Including an absorption step (such as passing digesta over Caco-2 cells in transwell inserts) between *in vitro* digestion and fermentation could overcome this. Another potential solution to this is to use dialysis to remove sugar and other water-soluble metabolites from the digesta ^42^, though this would result in loss of water soluble (but poorly absorbed) phenolics like CGA. As a result of not modeling absorption, the observed inhibition of TMA-d_9_ may be due to the presence of compounds that would not normally be in the colon, rather than phenolics, which have generally low bioavailability and would be present during fermentation. We observed this issue in experiments with whole blueberries, pulp, and sugar. However, this was addressed as much as possible by removing the pulp, which contains most of the substrates not found in the colon, and focusing on inhibition from the skin. We elected to use fiber and phenolic-rich skin for our multi-genotype experiment, as this material is physiologically relevant to the colon and reduces interferences.

There are inherent limitations to ex *vivo* fecal fermentations. Fecal samples were sourced from healthy donors, and their TMA production capacity compared to donors with CVD-associated dysbiosis is unknown. TMA production depends on bacteria that produce TMA lyase. By incorporating fecal samples from subjects with CVD, it may be possible to increase the abundance of bacteria that metabolize choline, creating an environment that could be more reflective of conditions seen in a state of CVD-associated dysbiosis. TMA production from known high TMA-producing donors may better reflect the conditions of the population of interest, as they likely include greater abundances of bacteria known to express cut*C/D* ^24,43–47^. Additionally, fecal samples are kept frozen and reanimated before fermentation. However, some bacteria may not grow outside the colon or survive the freeze-thaw and reanimation process, although a cryoprotectant is added to the samples. Furthermore, fecal samples are not fully representative of all regions of the colon. The use of pooled fecal samples is designed to create a more representative sample. This approach is beneficial for introducing bacteria from individuals with diverse lifestyles, living environments, and diets. However, it does not allow for an in-depth examination of the individual variations in choline metabolism to TMA. An inter-individual study could be conducted by keeping the OpenBiome fecal samples separate, to allow for comparison of choline conversion to TMA across different individuals.

Animal models would help address limitations associated with *in vitro* models. *In vitro* models tend to represent acute conditions, capturing short-term changes in enzyme production rather than shifts in the microbiome, due to the brief observation period. Models like MiGut, SHIME, and TIM-2 represent longer time frames *in vitro* and provide a more faithful simulation of the gut environment. However, they are more cumbersome, costly, and have lower throughput compared to our model ^48–50^. Longer *in vivo* studies would capture microbial shifts over time.

The blueberries themselves posed several challenges. There is limited information in the literature regarding blueberry fiber, making it difficult to determine exact ratios of the types of fibers. To address this, fiber extraction from blueberries would be necessary; however, this is time-consuming and outside the scope of this project. Removing the skin from the blueberries was challenging due to their small and delicate nature. Frozen blueberries were particularly difficult to blanch and separate the peel from the pulp without issues. Peeling by hand was a time-intensive process that increased the risk of contamination, either by exposing the berries to non-frozen temperatures for extended periods or by accidentally leaving peel in on the pulp.

The CGA content was measured for the whole blueberry across 20 different highbush genotypes, rather than just the skin, due to time limitations and the lengthy process of peeling the berries for the extracts used in SPE and LC-MS analysis. While previous experiments have shown that the majority of phenolics and CGA are contained in the peel, this was not specifically confirmed for the peels used in the final experiment. As a result, it is possible that the lack of differentiation in the TMA-d_9_ inhibition could be due to the smaller variations in the peel across all genotypes, compared to the greater differences observed in the whole fruit. The variations in the fruit could be attributed to the amount of peel rather than its concentration of CGA. Fruit sizes varied between genotypes, with some being much smaller with higher peel to pulp ratios in smaller berries. This was not accounted for in the experiment, as the amount of peel used was the same for all genotypes. As a result, the amounts may not accurately reflect the actual serving size of whole berries for each genotype, as each fruit had a different peel-to-pulp ratio.

## 5. Conclusions

Our data suggest that although highbush blueberry skin inhibits the production of pro-atherogenic TMA by human gut bacteria, CGA does not drive this activity. Given the lack of available interventions to blunt TMAO production, the present data showing that blueberry skins at translatable human doses inhibit the key step in this process are provocative. Future studies are needed to elucidate the mechanism of inhibition, identify the compound(s) responsible, and assess interindividual variability in efficacy. Given the fact that blueberries are widely available and consumed, the translatability of this effect to humans is promising. Pilot clinical studies should be designed in individuals exhibiting a high TMAO production phenotype.

## Author contributions

**Ashley M. McAmis:** methodology, validation, formal analysis, investigation, data curation, writing - original draft, writing - review & editing, visualization; **Michael G. Sweet:** methodology, formal analysis, investigation, writing - review & editing, visualization; **Sydney Chadwick-Corbin:** methodology, investigation, writing - review & editing; **Juanita G. Ratliff:** investigation, writing - review & editing; **Molla Fentie Mengist:** methodology, writing - review & editing; **Nahla V. Bassil:** resources, writing - review & editing; **Pon Velayutham Anandh Babu:** conceptualization, methodology, writing - review & editing, funding acquisition; **Massimo Iorizzo:** conceptualization, methodology, resources, writing - review & editing, funding acquisition; **Andrew P. Neilson:** conceptualization, methodology, formal analysis, data curation, writing - review & editing, visualization, supervision, project administration, funding acquisition.

## Data availability

The data supporting this article have been presented in the figures and tables (including supplementary figures and tables). Raw data are available upon request from the authors.

## Acknowledgements

The authors wish to acknowledge the contributions of Dr. Keith Harris (Department of Food, Bioprocessing, and Nutrition Sciences, North Carolina State University, Raleigh, NC) who suggested the use of an ATP-based microbial detection/viability assay. This work was supported by the Agriculture and Food Research Initiative Foundational and Applied Science Program, project award no. 2024-67017-42462, from the U.S. Department of Agriculture’s National Institute of Food and Agriculture. AMM was supported by funds from the Plants for Human Health Institute (North Carolina State University, Kannapolis, NC). MI was also supported by the United States Department of Agriculture National Institute of Food and Agriculture (USDA-NIFA), Hatch project 1008691. Support for APN was provided by the North Carolina Agricultural Research Service (NCARS) and the Hatch Program (project 7005424) of the National Institute of Food and Agriculture (NIFA), U.S. Department of Agriculture. We also acknowledge the staff of the USDA ARS NCGR for care and maintenance of blueberry genetic resources under CRIS Project 2072-21000-059-000D.

## References

1 Cardiovascular diseases (CVDs), https://www.who.int/news-room/fact-sheets/detail/cardiovascular-diseases-(cvds), (accessed January 30, 2024).

2 What is Cardiovascular Disease?, https://www.heart.org/en/health-topics/consumer-healthcare/what-is-cardiovascular-disease, (accessed January 30, 2024).

3 Atherosclerosis - What Is Atherosclerosis?, https://www.nhlbi.nih.gov/health/atherosclerosis, (accessed January 30, 2024).

4 N. Yoshida, T. Yamashita and K. Hirata, Diseases, 2018, 6, 56.

5 Y. Zhu, Q. Li and H. Jiang, APMIS, 2020, 128, 353–366.

6 L. Iglesias-Carres, M. D. Hughes, C. N. Steele, M. A. Ponder, K. P. Davy and A. P. Neilson, The Journal of Nutritional Biochemistry, 2021, 91, 108600.

7 R. A. Koeth, Z. Wang, B. S. Levison, J. A. Buffa, E. Org, B. T. Sheehy, E. B. Britt, X. Fu, Y. Wu, L. Li, J. D. Smith, J. A. DiDonato, J. Chen, H. Li, G. D. Wu, J. D. Lewis, M. Warrier, J. M. Brown, R. M. Krauss, W. H. W. Tang, F. D. Bushman, A. J. Lusis and S. L. Hazen, Nat Med, 2013, 19, 576–585.

8 J. R. Ussher, G. D. Lopaschuk and A. Arduini, Atherosclerosis, 2013, 231, 456–461.

9 S. H. Zeisel and K.-A. da Costa, Nutrition Reviews, 2009, 67, 615–623.

10 Anatomy & Function of Your Portal Vein, https://my.clevelandclinic.org/health/body/25048-portal-vein, (accessed March 2, 2024).

11 X. Li, J. Hong, Y. Wang, M. Pei, L. Wang and Z. Gong, Frontiers in Molecular Biosciences, DOI:10.3389/fmolb.2021.733507.

12 L. Jing, H. Zhang, Q. Xiang, L. Shen, X. Guo, C. Zhai and H. Hu, Front Cardiovasc Med, 2022, 9, 864600.

13 A. A. Amara and A. Shibl, Saudi Pharm J, 2015, 23, 107–114.

14 H. Bodke and S. Jogdand, Cureus, 14, e31313.

15 Z. Wang, A. B. Roberts, J. A. Buffa, B. S. Levison, W. Zhu, E. Org, X. Gu, Y. Huang, M. Zamanian-Daryoush and M. K. Culley, Cell, 2015, 163, 1585–1595.

16 Flavin monooxygenase 3, the host hepatic enzyme in the metaorganismal trimethylamine N-oxide-generating pathway, modulates platelet responsiveness and thrombosis risk - ScienceDirect, https://www.sciencedirect.com/science/article/pii/S1538783622023698, (accessed March 27, 2024).

17 Trimethylaminuria (’fish odour syndrome’), https://www.nhs.uk/conditions/trimethylaminuria/, (accessed March 25, 2024).

18 J. Messenger, S. Clark, S. Massick and M. Bechtel, J Clin Aesthet Dermatol, 2013, 6, 45–48.

19 A. C. Schmidt and J.-C. Leroux, Drug Discovery Today, 2020, 25, 1710–1717.

20 L. Iglesias-Carres, M. D. Hughes, C. N. Steele, M. A. Ponder, K. P. Davy and A. P. Neilson, The Journal of Nutritional Biochemistry, 2021, 91, 108600.

21 L. Iglesias-Carres, L. A. Essenmacher, K. C. Racine and A. P. Neilson, Nutrients, 2021, 13, 1466.

22 Bioaccessibility and bioavailability of phenolic compounds, https://www.sciopen.com/article/10.31665/JFB.2018.4162, (accessed January 18, 2025).

23 Are you absorbing the nutrients you eat?, https://www.canr.msu.edu/news/are_you_absorbing_the_nutrients_you_eat, (accessed January 13, 2025).

24 S. Rath, B. Heidrich, D. H. Pieper and M. Vital, Microbiome, 2017, 5, 54.

25 A. Rodriguez-Mateos, T. Cifuentes-Gomez, S. Tabatabaee, C. Lecras and J. P. E. Spencer, J Agric Food Chem, 2012, 60, 5772–5778.

26 V. Gavrilova, M. Kajdžanoska, V. Gjamovski and M. Stefova, J. Agric. Food Chem., 2011, 59, 4009–4018.

27 M. F. Mengist, M. H. Grace, J. Xiong, C. D. Kay, N. Bassil, K. Hummer, M. G. Ferruzzi, M. A. Lila and M. Iorizzo, Front Plant Sci, 2020, 11, 370.

28 A. K. Satheesh Babu, C. Petersen, L. Iglesias-Carres, H. A. Paz, U. D. Wankhade, A. P. Neilson and P. V. Anandh Babu, BioFactors, DOI:10.1002/biof.2014.

29 M. F. Mengist, M. H. Grace, T. Mackey, B. Munoz, B. Pucker, N. Bassil, C. Luby, M. Ferruzzi, M. A. Lila and M. Iorizzo, Frontiers in Plant Science.

30 L. Iglesias-Carres, K. C. Racine and A. P. Neilson, Food Funct., 2022, 13, 8022–8037.

31 L. Iglesias-Carres, A. Bruno, I. D’Antuono, V. Linsalata, A. Cardinali and A. P. Neilson, Journal of Functional Foods, 2023, 107, 105674.

32 Development of a genetic framework to improve the efficiency of bioactive delivery from blueberry - PubMed, https://pubmed.ncbi.nlm.nih.gov/33057109/, (accessed January 1, 2025).

33 In vitro evidences of the globe artichoke antioxidant, cardioprotective and neuroprotective effects - ScienceDirect, https://www.sciencedirect.com/science/article/pii/S1756464623002748?via%3Dihub, (accessed January 10, 2024).

34 L. Bresciani, M. Dall’Asta, C. Favari, L. Calani, D. D. Rio and F. Brighenti, Food Funct., 2018, 9, 6470–6483.

35 L. Iglesias-Carres, E. S. Krueger, J. A. Herring, J. S. Tessem and A. P. Neilson, J. Agric. Food Chem., 2022, 70, 3207–3218.

36 A. K. Satheesh Babu, C. Petersen, L. Iglesias-Carres, H. A. Paz, U. D. Wankhade, A. P. Neilson and P. V. Anandh Babu, BioFactors, 2024, 50, 392–404.

37 G. I. Sandle, Gut, 1998, 43, 294–299.

38 E. Jameson, M. Quareshy and Y. Chen, Methods, 2018, 149, 42–48.

39 R. A. Koeth, B. S. Levison, M. K. Culley, J. A. Buffa, Z. Wang, J. C. Gregory, E. Org, Y. Wu, L. Li and J. D. Smith, Cell metabolism, 2014, 20, 799–812.

40 P. Day-Walsh, E. Shehata, S. Saha, G. M. Savva, B. Nemeckova, J. Speranza, L. Kellingray, A. Narbad and P. A. Kroon, Eur J Nutr, 2021, 60, 3987–3999.

41 J. E. Haarhuis, P. Day-Walsh, E. Shehata, G. M. Savva, B. Peck, M. Philo and P. A. Kroon, Molecular Nutrition & Food Research, n/a, e70166.

42 Q. Yang, M. Van Haute, N. Korth, S. E. Sattler, J. Toy, D. J. Rose, J. C. Schnable and A. K. Benson, Nat Commun, 2022, 13, 5641.

43 J. Chen, B. Yu, D. Chen, P. Zheng, Y. Luo, Z. Huang, J. Luo, X. Mao, J. Yu and J. He, Appl Microbiol Biotechnol, 2019, 103, 8157–8168.

44 Z. Wang, K. Lam, J. Hu, S. Ge, A. Zhou, B. Zheng, S. Zeng and S. Lin, Food Sci Nutr, 2019, 7, 579–588.

45 Anaerobic choline metabolism in microcompartments promotes growth and swarming of Proteus mirabilis - PMC, https://www.ncbi.nlm.nih.gov/pmc/articles/PMC5026066/, (accessed March 28, 2024).

46 J. Song, N. Zhou, W. Ma, X. Gu, B. Chen, Y. Zeng, L. Yang and M. Zhou, Food Funct., 2019, 10, 2947–2957.

47 Chlorogenic Acid-Induced Gut Microbiota Improves Metabolic Endotoxemia - PMC, https://www.ncbi.nlm.nih.gov/pmc/articles/PMC8716487/, (accessed March 28, 2024).

48 T. Van de Wiele, P. Van den Abbeele, W. Ossieur, S. Possemiers and M. Marzorati, in The Impact of Food Bioactives on Health: in vitro and ex vivo models, eds. K. Verhoeckx, P. Cotter, I. López-Expósito, C. Kleiveland, T. Lea, A. Mackie, T. Requena, D. Swiatecka and H. Wichers, Springer, Cham (CH), 2015.

49 W. A. Davis Birch, I. B. Moura, D. J. Ewin, M. H. Wilcox, A. M. Buckley, P. R. Culmer and N. Kapur, Microb Biotechnol, 2023, 16, 1312–1324.

50 E. Maas, J. Penders and K. Venema, Journal of Fungi, 2023, 9, 104.

