## Supplementary Information for "Inhibition of pro-atherogenic trimethylamine production from choline by human gut bacteria is not determined by varying chlorogenic acid content in highbush blueberries"

for

**SUPPLEMENTARY METHODS**
(sections correspond to the main manuscript methods sections)

**2.1 Chemicals*:***

Fecal samples were obtained from OpenBiome (Cambridge, MA, USA). Urea (CAS# 57-13-6), uric acid (CAS# 69-93-2), KCl (CAS# 7447-40-7), NaCl (CAS# 7647-14-5), $NaHCO_3$ (CAS# 144-55-8), $Na_2HPO_4$ (CAS# 7558-79-4), PBS (CAS# 7447-40-7, 7647-145-5), formic acid (CAS# 64-18-6), methanol (CAS# 67-56-1), acetone (CAS# 67-64-1), acetonitrile (CAS# 75-05-8), concentrated ammonia (CAS#1336-21-6, 7732-18-5), $Na_3PO_4$ (CAS# 7601-54-9), $ZnSO_4$ (CAS# 7446-20-0) and sodium acetate (CAS# 6131-90-4) were obtained from Thermo Fisher Scientific (Hampton, NH). Mucin (porcine stomach, CAS# 84082-62-4), α-amylase (porcine pancreas, CAS# 0009000902), $Na_2SO_4$ (CAS# 7757-82-6), pepsin powder (porcine gastric mucosa, CAS# 9001-75-6). NaOH (CAS# 1310-73-2), pancreatin (porcine pancreas, CAS# 8049-47-6), lipase (porcine pancreas, CAS# 9001-62-1), bile extract (porcine, CAS# 8008-63-7), peptone water, yeast extract (CAS# 8013-01-2), $MgSO_4 \times 7H_2O$ (CAS# 10034-99-8), $CaCl_2 \times 6H_2O$ (CAS# 7774-34-7), hemin (CAS# 1600-13-5), bile salts, Tween 80 (CAS# 9005-65-6), vitamin K1 (CAS# 84-80-0), resazurin (CAS# 62758-13-8), L-cysteine (CAS# 52-90-4), choline- $d_9$ (CAS# 61037-86-3), ethyl bromoacetate (CAS# 105-36-2), sodium carbonate (CAS# 497-19-8), Folin-Ciocalteu reagent, ammonium formate (CAS# 540-69-2), Diaion HP-20 resin (CAS# 9052-95-3), TMA- $d_9$ (CAS# 18856-86-5), pectin from apple (CAS# 9000-69-5), cellulose (microcrystalline, 20 μm, CAS# 9004-34-6), D-(−)-Fructose (CAS# 57-48-7), and D-(+)-Glucose (CAS#50-99-7) were obtained from MilliporeSigma (Burlington, MA). Acetic acid (CAS# 64-19-7) and D-(+)-Sucrose (CAS# 57-50-1) were obtained from VWR (Radnor, PA). Choline-1-$^{13}C$-1,1,2,2, -$d_4$, choline-1,2-$^{13}C_2$, TMA-$^{13}C_3$-$^{15}N$, and TMA-$^{13}C_3$-$d_9$ were obtained from Cambridge Isotope Laboratories (Tewksbury, MA). Hemicellulose (CAS#9034-32-6) was obtained from BOC Sciences (Shirley, NY). $N_2$ gas was obtained from Airgas, Charlotte, NC, USA.

### General Methods

**2.3 General Gastrointestinal Digestion:**

50 mL Digestion. For a 50 mL oral digestion a ratio of 1 g of solid sample was mixed with 6 mL of an oral phase solution (0.4 g/L urea, 30 mg/L uric acid, 10.6 g/L α-amylase, 50 mg/L mucin, 1.792 g/L KCl, 1.776 g/L $Na_3PO_4$, 1.140 g/L $Na_2SO_4$, 0.596 g/L NaCl, 3.388 g/L $NaHCO_3$) homogenized in a 50 mL falcon tube and blanketed with $N_2$ gas, capped, and placed in an incubator at 37°C, and shaken at 120 opm (oscillations per minute) for 10 minutes. The gastric digestion occurred after the 10 minutes of shaking. The tubes were removed from the incubator and immediately placed on ice, brought to a volume of 30 mL using saline (0.9% NaCl in water), 2 mL of pepsin solution (20 mg/mL in HCL 0.1M) was added. Adjustment of pH to 2.5 ± 0.1 was achieved using 1M HCl and a pH probe. The volume was brought to 40 mL using saline. The tube was blanketed with $N_2$ gas, capped, placed in an incubator at 37°C, and shaken at 120 opm for 1 hour. The intestinal digestion occurred after shaking for 1 hour. The tubes were removed from the incubator and immediately placed on ice. Adjustment of pH to 6.5 ± 0.1 was achieved using 1M $NaHCO_3$. Then 2 mL of pancreatin (20 mg/mL in 0.1 NaHCO3)/ lipase (10 mg/mL in 0.1 NaHCO3) mixture was added. 3 mL of bile extract (30 mg/ mL in 0.1 $NaHCO_3$) was added. The final volume was brought to 50 mL with saline, blanketed with $N_2$ gas, capped, and placed in an incubator at 37°C, and shaken at 120 opm for 2 hours. The crude digesta was utilized. 5 mL of the same treatments were pooled in a new tube and stored in a −80°C freezer. The frozen digesta was lyophilized to remove water with solubilized oxygen and stored in a − 80°C freezer.

15 mL Digestion. For a 15 mL oral digestion, a ratio of 1 g of solid sample per 1.8 mL in oral phase solution (0.4 mg/mL urea, 0.03 mg/mL uric acid, 10 g mg/mL α-amylase per g food digested, 0.05 mg/mL mucin, 1.792 g/L KCl, 1.776 g/L $Na_3PO_4$, 1.140 g/L $Na_2SO_4$, 0.596 g/L NaCl, 3.388 g/L $NaHCO_3$), homogenized in a 15 mL falcon tube and blanketed with $N_2$ gas, capped, placed in an incubator at 37°C, and shaken at 120 rpm for 10 minutes. The gastric digestion occurred after the 10 minutes of shaking. The tubes were removed from the incubator and immediately placed on ice, brought to a volume of 9.6 mL using saline (0.9% NaCl in water), 0.6 mL of pepsin solution (20 mg/mL in HCL 0.1M) was added. Adjustment of pH to 2.5 ± 0.1 was achieved using 1M HCl and a pH probe. The volume was brought to 12 mL using saline. The tube was blanked with $N_2$ gas, capped, placed in an incubator at 37°C, and shaken at 120 rpm for 1 hour. The intestinal digestion occurred after shaking for 1 hour. The tubes were removed from the incubator and immediately placed on ice. Adjustment of pH to 6.5 ± 0.1 was achieved using 1M $NaHCO_3$. Then 0.6 mL of pancreatin (20 mg/mL in 0.1 NaHCO3)/ lipase (10 mg/mL in 0.1 NaHCO3) mixture was added. 0.9 mL of bile extract (30 mg/ mL in 0.1 $NaHCO_3$) was added. The final volume was brought to 15 mL with saline, blanked with $N_2$ gas, capped, and placed in an incubator at 37°C, and shaken at 120 rpm for 2 hours. The crude digesta was utilized. 5 mL the same treatments was pooled in a new tube and stored at −80°C. The frozen digesta was lyophilized to remove water with solubilized oxygen and stored in a −80°C freezer.

A "blank" or "control" digesta was made with saline in place of food material and treated the same as the other treatments.

**2.4 Growth Media Preparation:** A final volume of 500 mL growth media was composed of 1 g peptone water, 1 g yeast extract, 50 mg NaCl, 20 mg $Na_2HPO_4$, 20 mg $KH_2PO_4$, 5 mL $MgSO_4 \times 7H_2O$ + $CaCl_2 \times 6H_2O$, 1 g $NaHCO_3$, 25 mg hemin, 250 mg bile salts, 1 mL Tween 80, 5 μL vitamin K1, 2 mL resazurin, and 250 mg L-cysteine. To prepare the 500 mL solution, two separate solutions at 250 mL each were prepared at 2X final concentration. Solution one included all components of growth media except for resazurin and L-cysteine. Solution two only contained resazurin and L-cysteine. The pH of both solutions was adjusted to 6.8 then separately filtered through a 0.22 µm sterile filter. Solution two was boiled for 10 minutes until clear. Both solutions were sparged to remove oxygen. The bottles containing the media were sealed with a septa disk, which allowed needles to pass $N_2$ (g) through without exposing the contents to ambient air. For proper gas exchange, the number of inlet needles (introducing $N_2$) is equal to the number of outlet needles. The bottles were continuously stirred at 250 rpm while $N_2$ is passed through. This process was maintained for a minimum of 8 hrs to effectively remove oxygen. After sparging, solutions were combined 1:1 in the anaerobic chamber ($O_2 \leq 10$ ppm) to a final volume of 500 mL at 1X concentration.

**2.5 Fecal Slurry preparation:** Fecal samples collected from healthy donors and that are screened for infectious diseases were obtained through OpenBiome (Cambridge, MA, USA) and stored immediately upon arrival in a −80°C freezer. OpenBiome provides information for all health histories including clinical data, pathogen screen results, and 16S rDNA sequences. Samples are prepared by OpenBiome in sterile 12.5% glycerol and o.9% saline buffer (2.5 mL of buffer per one gram of stool) then filtered through a 330 µm filter and frozen at −80°C. The fecal slurries were prepped 12 hrs before the start of the fermentation. Fecal slurry volume varied based on individual experimental parameters. At least two different fecal samples were combined with at least 16 mL of growth media in a 50 mL tube and vortexed until homogenized. The fecal slurry was left uncapped in the anaerobic chamber for at least 12 hrs. The slurry was filtered immediately before the start of the fermentation. The remaining 2 mL of media was used to rinse residual fecal slurry through the filter to bring to final volume.

**2.6 General Fermentation Conditions:** Fermentations took place inside a 4-glove 855-ACB anaerobic chamber (Plas-Labs, Lansing, MI, USA). The chamber was shifted to anaerobic conditions 24 hrs before the start of the fermentation. This was done by sanitizing the chamber, and all equipment inside the chamber, using 70% ethanol and left to dry. The chamber was then purged of oxygen using a mixed gas and $N_2$. A day before the fermentation was performed, ≥10 gas exchanges first with $N_2$ (Airgas, Charlotte, NC, USA) then mixed gas containing 5% $H_2$, 5% $CO_2$ and 90% $N_2$ (ARC3, Raleigh, NC, USA) until chamber conditions reached $H_2$ (2-3%) and $O_2$ levels below 20 ppm monitored with a CAM-12 anaerobic monitor (Coy Lab Products, Grass Lake, MI, USA). Once anaerobic conditions were achieved, the heater inside the chamber was turned on to 37°C, and a palladium catalyst was placed on top before leaving it to stabilize overnight. Conditions were monitored throughout the fermentation experiment.

**2.7 General Fermentation Procedure:** Fermentations were carried out as per methodology developed by Iglesias-Carres *et al.*[1]. The anaerobic chamber was prepped as described above. To control for endogenous choline and other TMA-lyase substrates leading to TMA metabolism in the absence of exogenous choline, choline-$d_9$ was added as a substrate at a final concentration of 100 µM to all treatments. Subsequent TMA-$d_9$ was measured to evaluate the effectiveness of the treatments. Lyophilized, digested samples (if any) were prepared as described in general gastrointestinal digestions and were reconstituted to 1X concentration of the initial volume before lyophilization using filter-sterilized, overnight-sparged PBS 1X inside anaerobic chamber. In 2 mL 96-well plates, 750 µL of growth media was mixed with 90 µL of choline-$d_9$ stock solution (2mM) in PBS 1X, 600 µL of reconstituted digesta samples, and 360 µL of fecal slurry (aliquots from at least two different donors pooled). The dilution of 600 μL of digesta into a final volume of 1800 μL is a conservative method, as it reflects a more diluted concentration, avoiding the potential overestimation of the inhibitory effect that might occur with less dilution of digesta. Additionally, one of the functions of the colon is to absorb water, which results in a more concentrated mixture of digestive enzymes and food in this region. As a result, higher concentrations of digesta may be present *in vivo*, potentially leading to greater inhibition. This method aims to mimic the physiological environment of the colon by accounting for the relatively dilute matrix containing intestinal fluids. Solutions were pre-heated at 37°C, and the start of fermentation (time 0 h) began when fecal slurry was inoculated into reaction mixture with the substrate. From 0 to 24 or 30 hours, 100 μL of sample was collected at various time points, combined with 100 μL of acetonitrile, and frozen at −80°C.

**2.8 Phenolic Fraction Extraction:**

Phenolic Fraction Preparation. A phenolic extraction was performed to generate a blueberry fraction that accurately represents phenolic compounds in an average serving of blueberries (~150 g) for use in a fermentation. To achieve this, an equal mixture of frozen (distributed by Food Lion, LLC Salisbury, NC 28147) and fresh (SBROCCO International INC. Mount Laurel, NJ 08054) whole blueberries were homogenized using a ninja bullet blender for 5 min. After homogenization, the mixture was weighed (246.83 g), aliquoted, spread thinly on three separate pans, and lyophilized until all water was removed. The resulting lyophilized material was weighed (43.40657 g) and frozen at −80°C.

Phenolic Fraction Extraction. For extraction, 26.007 g of lyophilized blueberry mixture representing a serving size of fresh weight blueberry (~150 g) was combined with 234 mL of extraction mixture containing acetone, water, and acetic acid (70:29.5:0.5). A total of 2 L of extraction mixture was made for a total of 8 extractions. The mixture of solvent and freeze-dried material was vortexed, blended for 1 min using a polytron (VWR 200), sonicated for 5 min in a water bath at 50 °C, and then centrifuged at 3248 x g for 10 min. The steps were repeated seven times until the supernatant was clear. Due to the soft, fibrous nature of the blueberry pellet, completely removing the supernatant proved challenging. As a result, approximately 90% of the supernatant was carefully removed without disturbing the pellet after each extraction. A total of 8 extractions were performed, and the remaining supernatant did not affect the extract material, as multiple extraction repeats included the leftover supernatant. About 1600 mL of the supernatant was collected from the extractions, pooled, and concentrated using a rotary evaporator at 45 °C under partial vacuum. After removing the material, the flask was rinsed twice with 25 mL of Milli-Q water to remove leftover extract material. The rinses were pooled with the extract (~600 mL total). The extract material was frozen at −80°C then lyophilized until all water was removed.

The resulting lyophilized extract material was weighed (23.05 g) and resuspended in ~ 200 mL acidified mill-Q water (0.1% formic acid). 607.43 g of dry, Diaion HP-20 resin was transferred to a beaker and covered with methanol by approximately 2", stirred, and left to sit for 15 minutes. After the 15 minutes, the methanol was decanted and the resin was covered with milli-Q water by approximately 2", stirred, and left to sit for 10 minutes. Enough milli-Q water was added to an empty 24/40 column to reach a 1" height. The resin was slowly poured into the column, as the excess water slowly drained out. Once all the resin was added, five 9 cm filter papers were added followed by a 1" layer of sand. The resin was equilibrated by adding 10 column volumes of milli-Q water, and the water was discarded. The resuspended extract material was added slowly to form a uniform layer on top of the sand layer. Another 1" layer of sand was added on top of the extract material. Four liters of acidified milli-Q water (0.1% formic acid) was added to the column, and the resulting eluent was collected in large bottles. Once all acidified water was collected; three liters of acidified methanol (0.1% formic acid) was added to the column, and the resulting eluent was collected in large bottles. Lastly, three liters of acidified acetone (0.1% formic acid) was added to the column, and the resulting eluent was pooled with the methanol fractions. The pooled methanol and acetone fractions were transferred into a round bottom flask and concentrated using a rotary evaporator under partial vacuum starting at 80°C for the methanol fractions and then lowering the temperature to 45°C for the acetone fractions. After removing the concentrated material, the flask was rinsed and sonicated twice with 50 mL milli-Q water and once with 25 mL of milli-Q water. The rinsed fractions were pooled with the extract material (~170 mL total), frozen at −80°C, and lyophilized to remove water. The final lyophilized extract was weighed (1.01251 g) and stored at −80°.

**2.9 Plant Phenolic Extraction and Analysis:** Plant-derived phenolic extract can be used in phenolic assays to measure relative amounts of phenolics in blueberry fractions. 250 mg of blueberry material was mixed with 2.5 mL of solvent A (80% methanol 2% formic acid in water). The samples were vortexed for 30 seconds, sonicated for 20 min, and centrifuged for 4 minutes at 3428 x g. The supernatant was removed and dried under nitrogen. 2.5 mL of solvent B (98%methanol 2% formic acid) was then added to all samples. The samples were vortexed for 30 seconds, sonicated for 20 min, and centrifuged for 4 min at 3428 x g. The supernatant was removed and combined with the previous supernatant. Steps for solvent B were repeated. Once supernatant was dried, the samples were stored at −80°. To resolubilize the dried supernatant, 3 mL of 0.1% formic acid in water was mixed into each sample. The samples were vortexed until homogenized and then filtered. Resolubilized extracts were stored at −80°.

<u>Phenolic Analysis: Folin- Ciocalteu Assay.</u> To quantify the total phenolics in the plant extracts, prepared as described in "*Plant Phenolic Extraction"*, a Folin assay was carried out in a 96 well plate. Each extract underwent a series of dilutions. Starting with the pure extract then diluting 10x, then diluting 100x, etc. A standard curve was made with a 2mg/mL gallic acid mixture with a series of 23- 2/3rds dilutions with the last dilution as pure water to represent a "blank". 20 µL of both the standard and extract samples were plated in triplicate. 140 µL of Folin reagent (2mL of Folin-Ciocalteu reagent mixed in 18 mL water) was mixed with all samples. The plate was covered with foil and protected from light for 30 min. After the 30 min, 100 uL of stop solution (25% sodium carbonate in water) was added, and the absorbance was taken at 765 nm. Total phenolics are expressed as mg of gallic acid equivalent (GAE)/ g of whole blueberry.

<u>SPE and LC-MS Analysis.</u> To quantify the relative amounts of CGA in plant material, a solid phase elution (SPE) followed by analysis using LC-MS was utilized following methodology described by Mengist *et al.*[2]. Plant extracts described in "*Plant Phenolic Extraction"* underwent a series of dilutions. Starting with the pure extract then diluting 10x, then diluting 100x, etc. A standard of 0.2 mM ethyl gallate was used to determine loss during the SPE process. An internal standard of 0.2 mM Taxifolin was used for the LC/MS. A standard curve was made with a

1mg/mL CGA mixture with a series of dilutions. Samples were plated in triplicate. 500 µL of sample and standard curve were mixed with 500 µL of 1% formic acid in water. A waste collection plate was placed on top of the StrataX filter plate (Phenomenex, Strata-X 96well plate 10mg/ well, Cat # 8E-S100-AGB) and was conditioned with 1 mL 1% formic acid in methanol, placed on a positive pressure manifold, and methanol was pulled through. The waste plate was emptied before 1 mL of 1% formic acid in water was pulled through using the positive pressure manifold. The waste plate was emptied before 1 mL of the sample mixed with 1% formic acid and 20 µL of Ethyl Gallate were plated and pulled through. Once the sample was pulled through, the filter was washed with 1 mL 1% formic acid in water twice and then washed with pure water. The waste plate was emptied, and samples were dried. The dried samples were resolubilized using 300 µL of 1% formic acid in water and left to sit for 10 min. After the 10 min, the samples were slowly pulled through the filter. 20 µL of Taxifolin was plated on a new 96-well plate with 160 µL of the final sample. Samples were then characterized on LC-MS. Separation was accomplished using a Waters Acquity UPLC system (Milford, MA, USA) with an ACQUITY UPLC BEH C18 column (1.7 µm, 2.1 mm x150 nm). Mobile phases include 0.1% formic acid in acetonitrile (A) and 2% formic acid in water (B) and phenolics were separated using a gradient elution method which involved a 6-minute run time, started with 100% solvent B for the first 30 seconds. Over the next 5 minutes, the solvent composition gradually shifted to 35% solvent A and 65% solvent B. In the final minute of the elution, the solvent composition shifted back to 100% solvent B. Flow rate was 0.5 mL/min, column temperature was set to 30°C, and autosampler was set to 4°C. The above system was coupled with a Waters Acquity triple quadrupole mass spectrometer. Source temperatures was 150°C. Capillary voltage was 2 kV, desolvation ($N_2$) and gas flow ($N_2$) were set to 600°C and 650 L/h, respectively. Electrospray ionization (ESI) was operated in negative mode. Data were collected using multiple reaction monitoring (MRM) in MS/MS mode (Table 3.1). MS/MS parameters for chlorogenic acid detection as presented above in **Supplementary Table 1**. A representative UPLC-MS/MS chromatogram of CGA is shown in.

**Supplementary Table 1.** Analyte multiple reaction monitoring (MRM) values for chlorogenic acid

| Analyte | MRM | Collision Energy (V) | Cone Voltage (V) |
|---|---|---|---|
| chlorogenic acid | 353.0228→84.9980 | 40 | 26 |

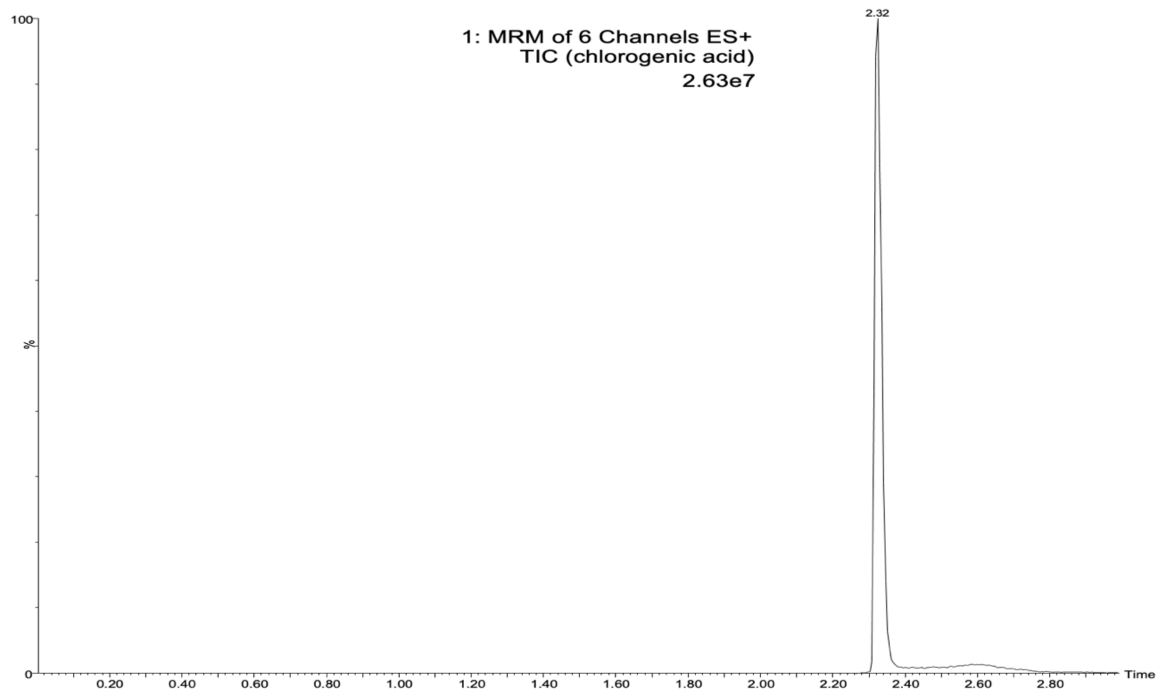


***Supplementary Figure 1.*** *Representative UPLC-MS/MS chromatogram of chlorogenic acid.*

**2.10 Digesta and Fermenta CGA Analysis:** To further identify the source of inhibition from the genetically diverse blueberry treatment, CGA analysis of the digesta and fermenta samples was performed. 1 mL of digesta sample was combined with 1 mL of acidified ACN (0.1% FA in ACN) *n*=2, vortexed, sonicated for 5 min, centrifuged (10 min at 17,000 x g), and filtered through 0.2 micron PTFE filter. Fermenta samples were combined 1:1 with ACN after sample collection during fermentation. To acidify these samples 10 µL of 0.425% formic acid in ACN was mixed with 75 µL of fermenta sample. Samples were filtered through Strata filter plates using centrifugation (10 min at 3428 x g) before LC-MS analysis. Separation was accomplished using a Waters Acquity UPLC system (Milford, MA, USA) with an ACQUITY UPLC BEH C18 column (1.7 µm, 2.1 mm x150 nm). Mobile phases include 0.1% formic acid in acetonitrile (A) and 2% formic acid in water (B) and phenolics were separated using a gradient elution method which involved a 3-minute run time, started with 100% solvent B for the first 90 seconds. Over the next minute, the solvent composition gradually shifted to 95% solvent A and 5% solvent B. In the final 90 seconds of the elution, the solvent composition shifted back to 100% solvent B. Flow rate was 0.5 mL/min, column temperature was set to 30°C, and autosampler was set to 4°C. The above system was coupled with a Waters Acquity triple quadrupole mass spectrometer. Source temperatures was 150°C. Capillary voltage was 3 kV, desolvation ($N_2$) and gas flow ($N_2$) were set to 200°C and 650 L/h, respectively. Electrospray ionization (ESI) was operated in positive mode. MS/MS parameters for chlorogenic acid detection as presented above in **Supplementary Table 1**.

**2.11 *BacTiter-Glo ATP Assay:*** Bacterial viability was quantified by luminescence using a commercial BacTiter-Glo Microbial Cell Viability Assay as described in *BacTiter-Glo ATP*

*Assay.* Experiments were performed to validate the BacTiter-Glo ATP Assay for use in our conditions.

<u>Interference determination.</u> 1mM ATP in PBS 1X was prepared and ATP standard curves were made in varying mediums that represented a component of the final fermentation mixture: PBS, growth media, PBS+ choline-$d_9$, PBS +CGA (10mM, 1mM, 10 µM and 1 µM), PBS + blueberry digesta, PBS + blank digesta. 100 µL of the serial dilutions of ATP in the various mediums were plated *n*=3 on white 96-well opaque round-bottom plates. Each well was mixed with 100 uL of reconstituted BacTiter-Glo reagent at room temperature and placed in the plate reader (SpectraMax iD3, Molecular Devices, USA) for 5 min to rest. Luminesce was read (Shake: 10s, integration: 2000, read height: 0.50mm) and results recorded.

The results of the interference determination experiment are presented in **Supplementary Figure 2** and **Supplementary Table 2**. ATP standard curves prepared in varying mediums exhibited different luminescence signals. Standard curves prepared in media representing blueberry digesta and blank digesta yielded the lowest luminescence signals compared to other treatments, with blueberry digesta showing the greatest suppression, likely due to the blue color of the blueberry. This suggests that digestive enzymes, along with the presence of food and associated color, interfere with the luminescence signals. In contrast, ATP standard curves made with choline-$d_9$ displayed the highest luminescence among all treatments. PBS and media treatments, which constitute the largest proportion of the final fermentation mixture, exhibited high luminescence readings. This suggests that any potential reduction in luminescence signals in fermenta containing fecal bacteria is more likely due to interference from the digesta background (particularly blueberry digesta), rather than from PBS or the media background.

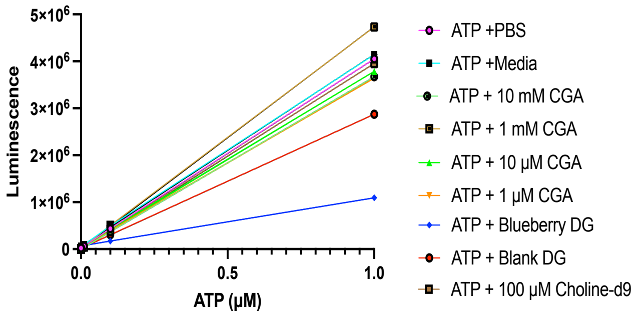


***Supplementary Figure 2.*** *ATP standard curves in varying mediums representing a component of the final fermentation mixture plated on a 96-well white round-bottom plate: chlorogenic acid (CGA), digesta (DG): PBS, growth media, 10 mM CGA, 1 mM CGA, 10 µM CGA, 1 µM CGA, blueberry DG, blank DG, and choline-$d_9$ (100 µM). Data represents mean SEM from n=3.*

**Supplementary Table 2.** Simple linear regression tabular results for ATP standard curves in varying mediums plated on white 96-well plates (*n*=3)

| | **Medium** | | | | | | | | |
|---|---|---|---|---|---|---|---|---|---|
| | | | **CGA** | | | | **Digesta** | | |
| **Regression value** | **PBS** | **Growth Media** | **10 mM** | **1 mM** | **10 µM** | **1 µM** | **Whole blueberry** | **Blank** | **Choline-$d_9$ (100 µM)** |
| $r^2$ | 1.000 | 0.9999 | 1.000 | 1.000 | 1.000 | 1.000 | 1.000 | 1.000 | 1000 |
| P-value (slope) | <0.0001 | <0.0001 | <0.0001 | <0.0001 | <0.0001 | <0.0001 | <0.000 | <0.0001 | <0.0001 |

<u>Cell washing validation.</u> Based on previous results which showed inhibited luminescence in treatments with digesta, particularly blueberry digesta, an experiment was conducted to evaluate whether "washing" fecal bacteria reduces fermentation background interferences. A fermentation was carried out in a 2 mL, 96-well plate following methodology described above in "*general fermentation procedures*". Treatments included growth media, growth media with fecal, blueberry digesta, blueberry digesta with fecal, blank digesta, and blank digesta with fecal. 100 uL of fermenta samples were plated in *n*=2 on a white 96-well opaque round-bottom plate at 12 h (Figure 4.3.) and 24 h (Figure 4.4.). The plate was centrifuged for 5 min at 4°C at 500 x g. The supernatant was removed from the first set of samples, and the pellet was resuspended in 100 µL of PBS 1X, this set of samples is referred to as "washed". The second set of samples were resuspended back into the original fermenta sample. All samples were mixed with 100 µL of reconstituted BacTiter-Glo reagent and placed in the plate reader for 5 min before reading the luminescence. Results were recorded and it was determined that samples are to be centrifuged and resuspended in 100 µL PBS 1X before adding 100 µL BacTiter-Glo reagent to reduce matrix interference and obtain a clear signal.

Results of the washing experiment are presented in **Supplementary Figure 3**. Treatments inoculated with fecal slurry exhibited a clear luminescence signal, providing evidence that the signal is originating from the fecal bacteria, rather than from other components of the fermentation matrix. Notably, the washed blueberry digesta and fecal treatment produced a luminescence signal that was twice as high as the same unwashed treatment, providing strong evidence that washing effectively reduces the background interferences from the digesta. At the 12-hour mark, a statistically significant difference was observed between the washed and unwashed samples that contained fecal bacteria, supporting the rationale for washing the fermenta samples to minimize background interference. At the 12 hr mark, treatments containing blueberry digesta exhibited significantly higher luminescence signals compared to all other treatments, suggesting blueberry digesta may promote bacterial growth. By the 24 hr mark, all treatments exhibited a luminescence signal, with W media + fecal and W blank + fecal exhibiting the highest signal. Interestingly, whole blueberry treatments at 24 hr displayed a signal even in the absence of fecal inoculation, which may be attributed to a potential bacterial contamination in those treatments. Overall, washing the fecal bacteria in the fermenta samples significantly reduced background interference, enhancing the detection of luminescence. This allowed the assay to specifically measure luminescence from the bacteria, rather without quenching interferences from the fermentation matrix. As a result, all fermentation treatments were "washed" in subsequent experiments

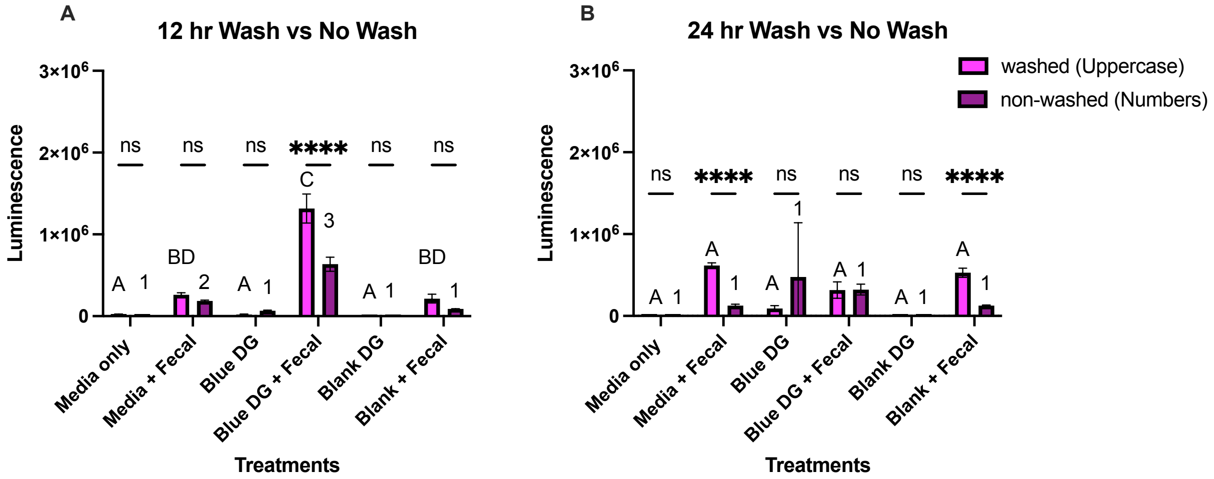


| | | | |
|---|---|---|---|
| Source of Variation | % of total variation | P value | P value summary |
| treatments x wash | 11.32 | <0.0001 | **** |
| treatments | 83.38 | <0.0001 | **** |
| wash | 3.564 | <0.0001 | **** |

| | | |
|---|---|---|
| Fixed effects (type III) | P value | P value summary |
| treatments | 0.2732 | ns |
| wash non wash | <0.0001 | **** |
| treatments x wash non wash | <0.0001 | **** |

***Supplementary Figure 3.*** *Luminescence signals of all treatments at the 12 hr (A) and 24 hr (B) mark. Washed (W), digesta (DG), and blue (whole blueberry digesta): media only, W media only, W media + fecal, media + fecal, W blue DG, Blue DG, W blue DG + fecal, blue DG + fecal, W blank DG, blank DG, W blank DG + fecal, blank DG + fecal. Different uppercase letters notate statistical differences between washed treatments (P<0.05) using a 2-way ANOVA and Sidak's multiple comparison test. Different numbers notate statistical differences between non-washed treatments (P<0.05) using Mixed-effect analysis and Sidak's multiple comparison test. Asterisks notate statistical differences between washed and non-washed within treatments (P<0.05) using a 2-way ANOVA and Tukey's multiple comparison test. Data represents mean SEM from n=2.*

Linearity of response. Next, we aimed to evaluate the impact of fecal slurry concentration on luminescence signal, a 5-fold diluted fecal slurry (1 mL of fecal + 4 mL of media) was prepped under anaerobic conditions 12 hrs prior to the start of the experiment. Six serial dilutions were then created by diluting the 5-fold fecal slurry to achieve 10-fold, 20-fold, 40-fold, 80-fold, and 160-fold concentrations by diluting 1:1 using the fecal suspension and media at each step. 100 µL from each dilution were plated *n*=3 on a white 96-well opaque round-bottom plate. The plate was centrifuged at 500 x g for 5 min, the supernatant was removed, and the pellet resuspended in PBS 1X. Following resuspension of the pellet, 100 µL of reconstituted BacTiter-Glo was mixed into the samples and left to sit in the dark for 5 min. Luminescence was read. Results from the relationship between cell concentration and luminesce are presented in **Supplementary Figure 4**, where luminescence signals for all dilutions are statistically different from one another. A linear trend is observed for the 10-fold, 20-fold, 40-fold, 80-fold, and 160-fold dilutions, with luminescence signals decreasing as fecal concentration becomes more diluted. However, the difference in luminescence signals between the 5-fold and 10-fold dilutions is less pronounced, suggesting that at very high and very low concentrations, differentiation becomes less linear. At higher concentrations, the luminescence signal levels off, rather than continuing to follow a linear pattern.

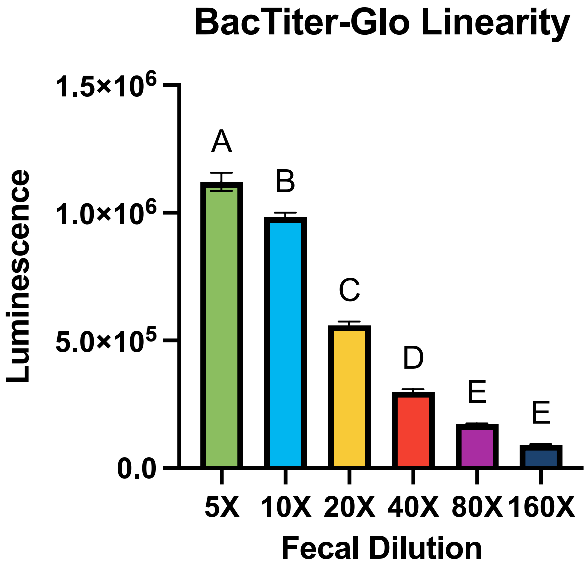


***Supplementary Figure 4.*** *BacTiter-Glo luminescence of varying concentrations of diluted fecal slurry: 5-fold, 10-fold, 20-fold, 40-fold, 80-fold, 160-fold. Different letters notate a statistical difference (P<0.05) using an ordinary one-way ANOVA with Tukey's multiple comparison test. Data represent mean ± SEM from n=3.*

**2.12 Extraction and Quantification of Choline-$d_9$ and TMA-$d_9$:** Sample extraction and quantification of choline-$d_9$ and TMA-$d_9$ from the stable isotope dilutions were carried out as described by Iglesias-Carres et al.[3]. Two choline internal standards are available for use, with only one utilized per experiment. Structures of choline internal standards are shown in **Supplementary Figure 5A-B**. After the sample collection, choline- $d_9$ is extracted in a 96-well plate. 25 µL of fermenta sample were mixed with 10 µL of $ZnSO_4$ solution, 100 µL acetonitrile, and 20 µL of either choline-1-$^{13}$C-1,1,2,2, -$d_4$ (Figure 3.1A.) or choline-1,2-$^{13}C_2$ (Figure 3.1 B.) solution (internal standard; 20µM). The plate was sonicated for 5 minutes in a water bath, and samples were then filtered through AcropreAdv 0.2 µm WWPTFE 96-well filtering plates (Pall Corporation, Port Washington, NY, USA) using centrifugation for 10 minutes at 3,400 x g. The reaction scheme for the experiment and analysis is shown in **Supplementary Figure 5C.** Samples are collected in a fresh 700 µL 96-well plate and frozen at −80°C. TMA- $d_9$ requires a derivatization process to the quaternary amine compound, ethyl betaine- $d_9$ (Figure 3.1C.). To analyze TMA- $d_9$, 25 µL of fermenta sample is mixed with 120 µL ethyl bromo acetate (20 mg/mL) solution, 9 µL of 28-30% ammonia, and 20 µL TMA-$^{13}C_3$-$^{15}$N (Figure 3.1D.) or TMA-$^{13}C_3$-$d_9$ (Figure 3.1E.) solution (internal standard; 20 µM, for derivatization to ethyl betaine-$^{13}C_3$-$^{15}$N or ethyl betaine-$d_9$, respectively). TMA internal standards and derivatization reactions are shown in **Supplementary Figure 5D-E.**

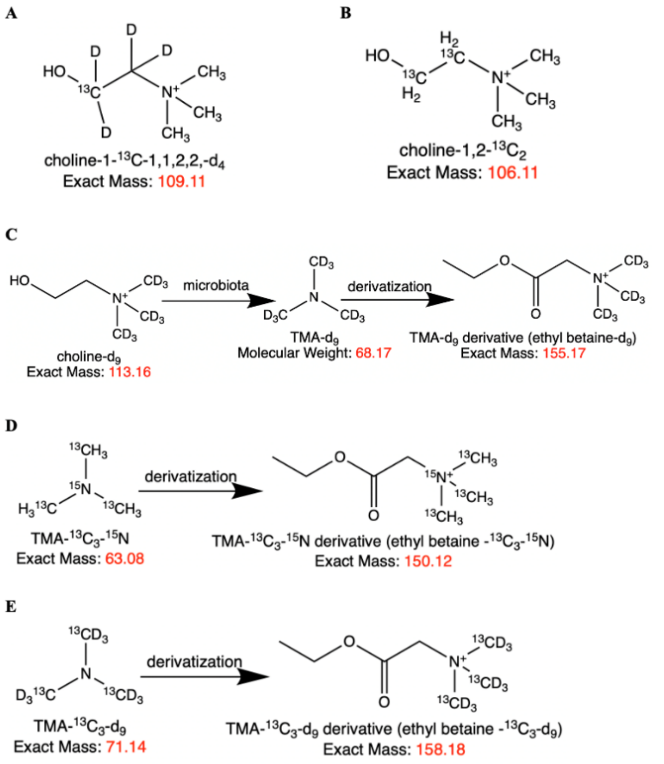


***Supplementary Figure 5**. Internal standards (A) choline-1-$^{13}C$-1,1,2,2, -$d_4$ and (B) choline-1,2-$^{13}C_2$. (C) experimental and analytical reaction scheme. (D) TMA-$^{13}C_3$-$^{15}N$ derivatization, TMA-$^{13}C_3$-$d_9$ derivatization.*

The samples are left to rest for 30 minutes at room temperature. After the 30 minutes, 120 µL 50% acetonitrile/0.025% formic acid in HPLC grade water is added to stop the reaction. TMA-$d_9$ samples are filtered as described for choline- $d_9$. After extraction, TMA- $d_9$ (Figure 3.3), TMA-$^{13}C_3$-$^{15}N$ (Figure 3.4) and TMA-$^{13}C_3$-$d_9$ (Figure 3.5) derivatized samples are analyzed using UPLC-ES-MS/MS. Choline-$d_9$ (Figure 3.6), choline-1,2-$_{13}C_2$ (Figure 3.7), and choline-1-$^{13}C$-1,1,2,2, -$d_4$ (Figure 3.8) were analyzed separately, but with the same method, from TMA-$d_9$, TMA-$^{13}C_3$-$d_9$,and TMA-$^{13}C_3$-$^{15}N$. Separation was accomplished using a Waters Acquity UPLC system (Milford, MA, USA) with an ACQUITY BEH HILIC column (1.7 µm, 2.1x100 nm) and an AQUITY BEH HILIC pre-column (1.7 µm, 2.1x4 nm). Mobile phases include 15 mM ammonium formate in water (pH 3.5) (A) and acetonitrile (B). Gradient is isocratic at 80% B for 3 minutes, flow rate 0.65 mL/min, column temperature set to 30°C, and autosampler set to 4°C. The above system is coupled with a Waters Acquity triple quadrupole mass spectrometer. Source and capillary temperatures are 150°C and 400°C, respectively. Capillary voltage is 0.60 kV, desolvation ($N_2$) and gas flow ($N_2$) set to 800 and 20 L/h, respectively. Electrospray ionization (ESI) operated in positive mode; data were collected using multiple reaction monitoring (MRM) in MS/MS mode (**Supplementary Table 3**). Standard curves are created by dividing choline-$d_9$ or TMA-$d_9$ values by their respective internal standard, then plotting those ratios against the known concentration of the standard analyte, to generate a slope equation. For fermentation samples, the choline-$d_9$ or TMA-$d_9$ values are divided by their respective internal standard and input into the slope equation to get calculate the actual concentration. These values are then multiplied by a factor of two to correct for dilution with acetonitrile. Finally, the values are plotted over time to produce kinetic curves for choline-$d_9$ utilization and TMA-$d_9$ production.

**Supplementary Table 3.** Analyte multiple reaction monitoring (MRM) values for choline and TMA pathway analysis

| Analyte | MRM | Collision Energy (V) | Cone Voltage (V) |
|---|---|---|---|
| TMA-$d_9$ | 155.33→127.21 | 34 | 20 |
| choline-$d_9$ | 113.32→69.08 | 40 | 16 |
| TMA-$^{13}C_3$-$^{15}N$ | 150.27→122.19 | 34 | 18 |
| TMA-$^{13}C_3$-$d_9$ | 158.20→130.13 | 34 | 16 |
| choline-1-$^{13}C$-1,1,2,2,-$d_4$ | 109.28→60.36 | 36 | 18 |
| choline-1,2-$^{13}C_2$ | 106.10→60.04 | 36 | 14 |

Representative chromatograms of all analytes are shown in **Supplementary Figures 6-11.**

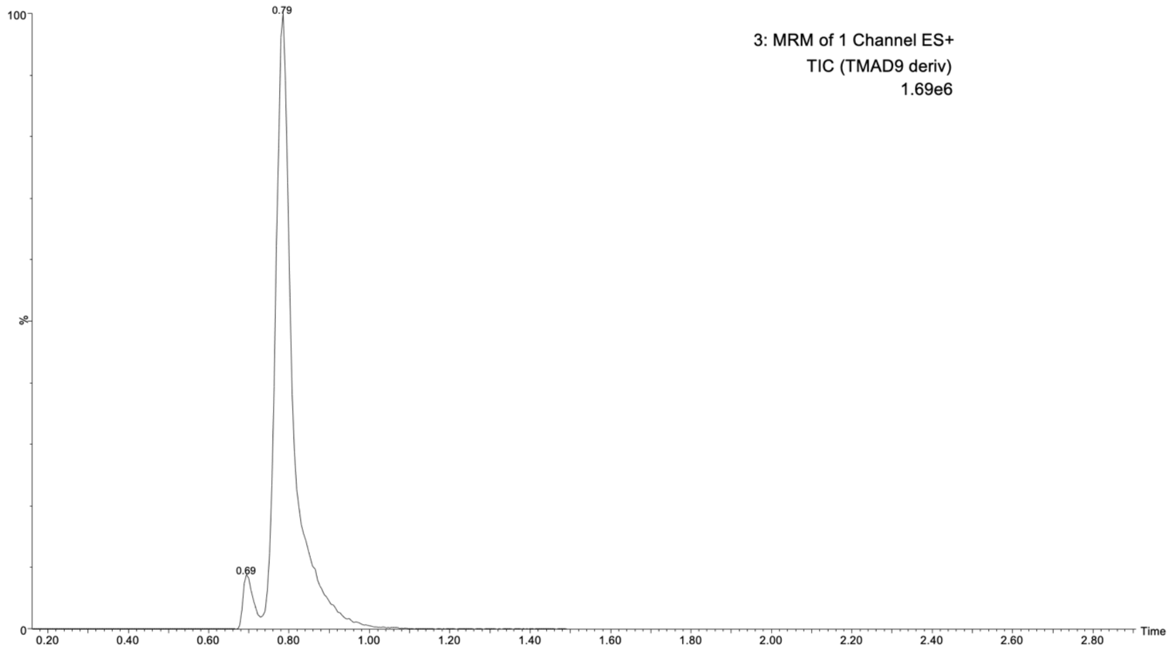


***Supplementary Figure 6***. *Representative UPLC-MS/MS chromatogram of derivatized TMA-$d_9$* (ethyl betaine-$d_9$).

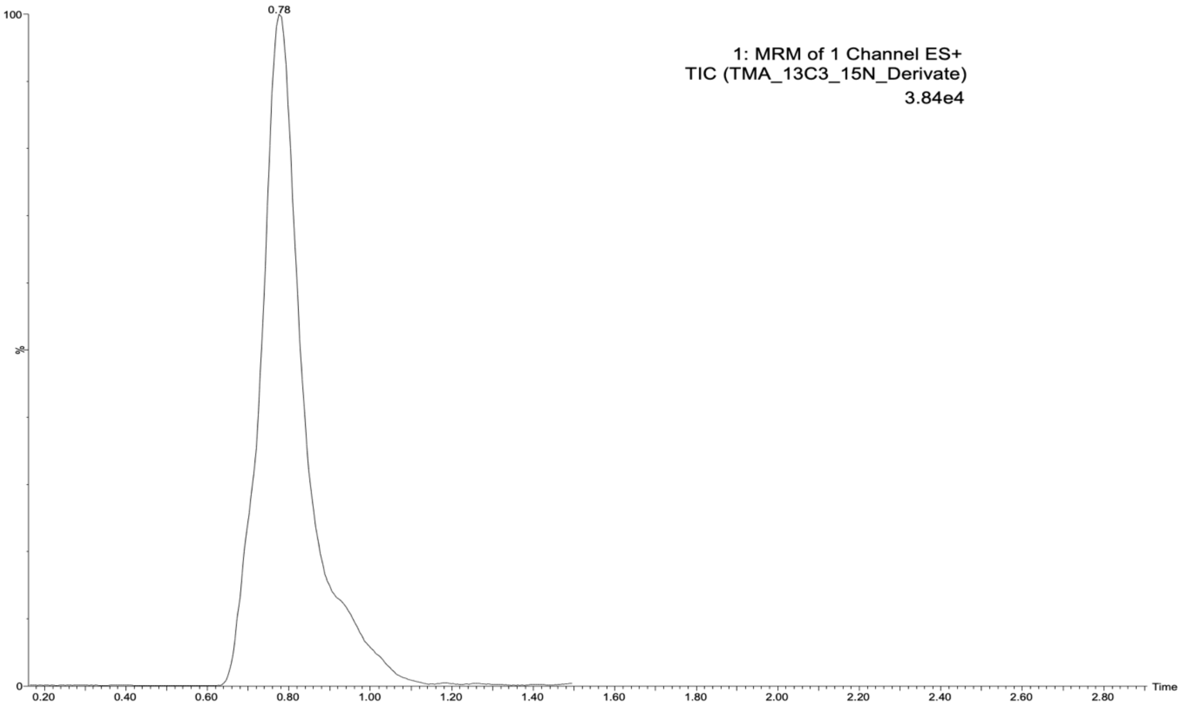


***Supplementary Figure** 7. Representative UPLC-MS/MS cof derivatized internal standard TMA-$^{13}C_3$-$^{15}N$ (ethyl betaine $^{13}C_3$-$^{15}N$).*

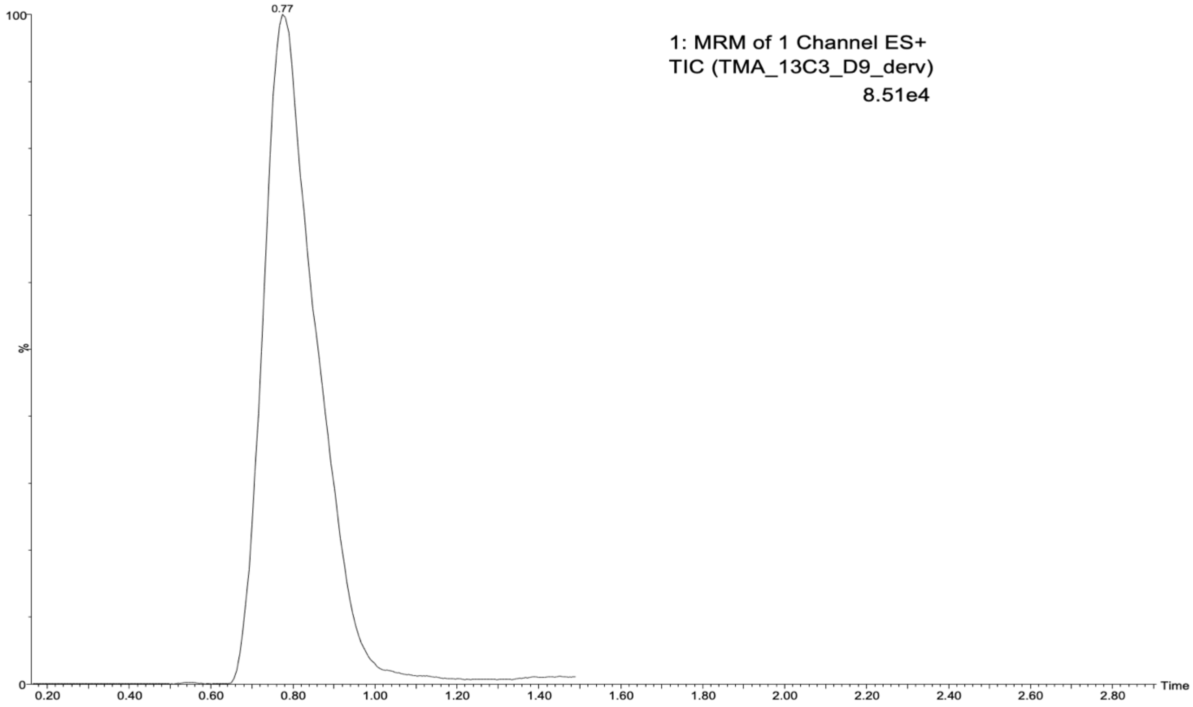


***Supplementary Figure 8**. Representative UPLC-MS/MS chromatogram of derivatized internal standard TMA-$^{13}C_3$-$d_9$ (ethyl betaine $^{13}C_3$- $d_9$).*

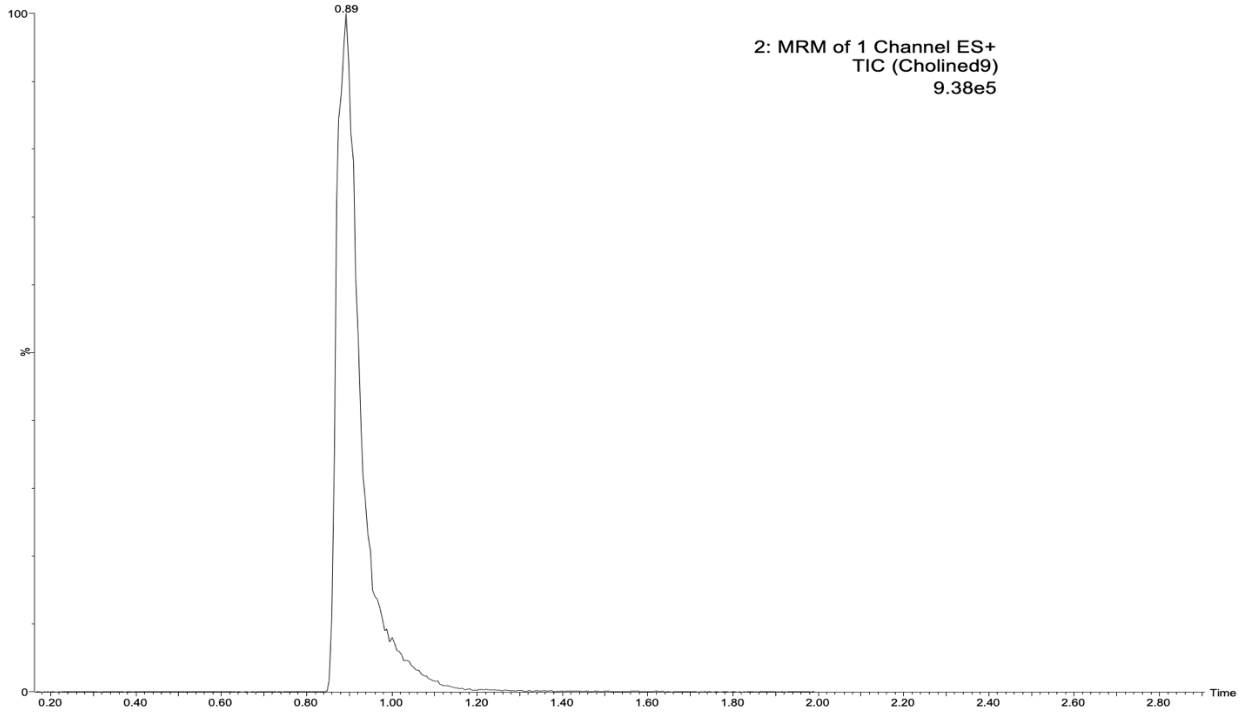


***Supplementary Figure 9***. *Representative UPLC-MS/MS chromatogram of choline-$d_9$.*

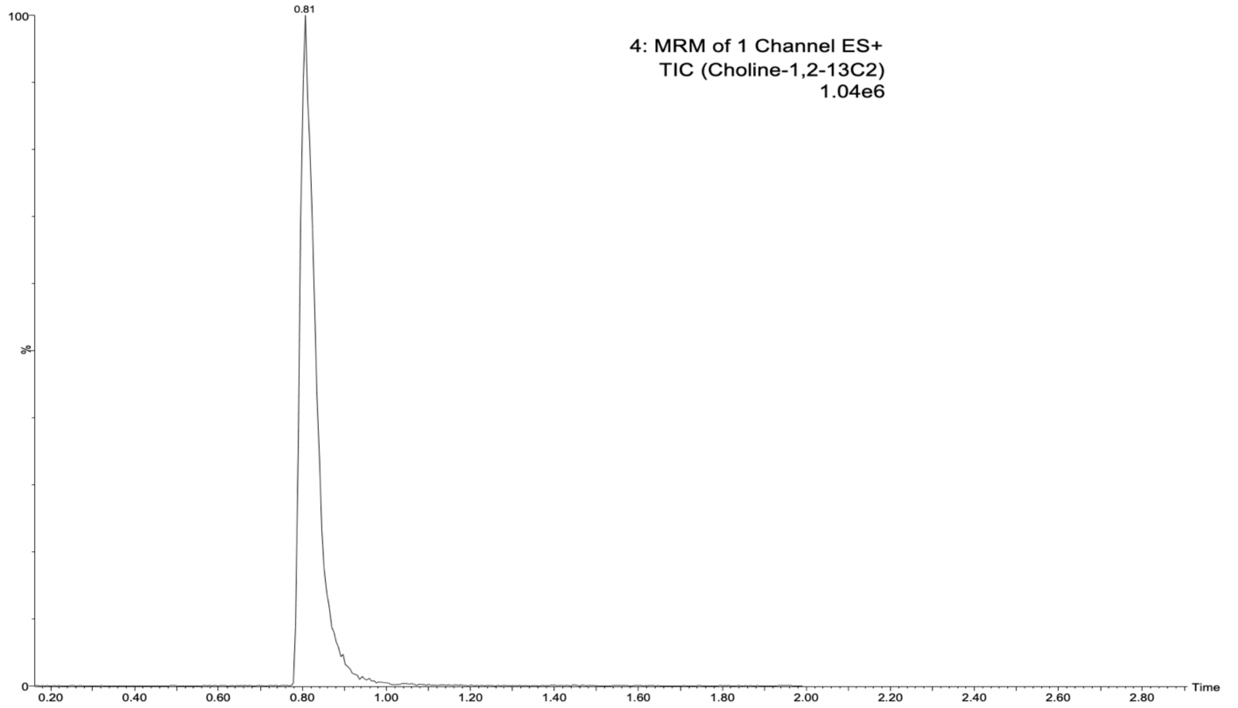


***Supplementary Figure 10***. *Representative UPLC-MS/MS chromatogram of internal standard choline-1-$^{13}$C-1,1,2,2,-$d_4$.*

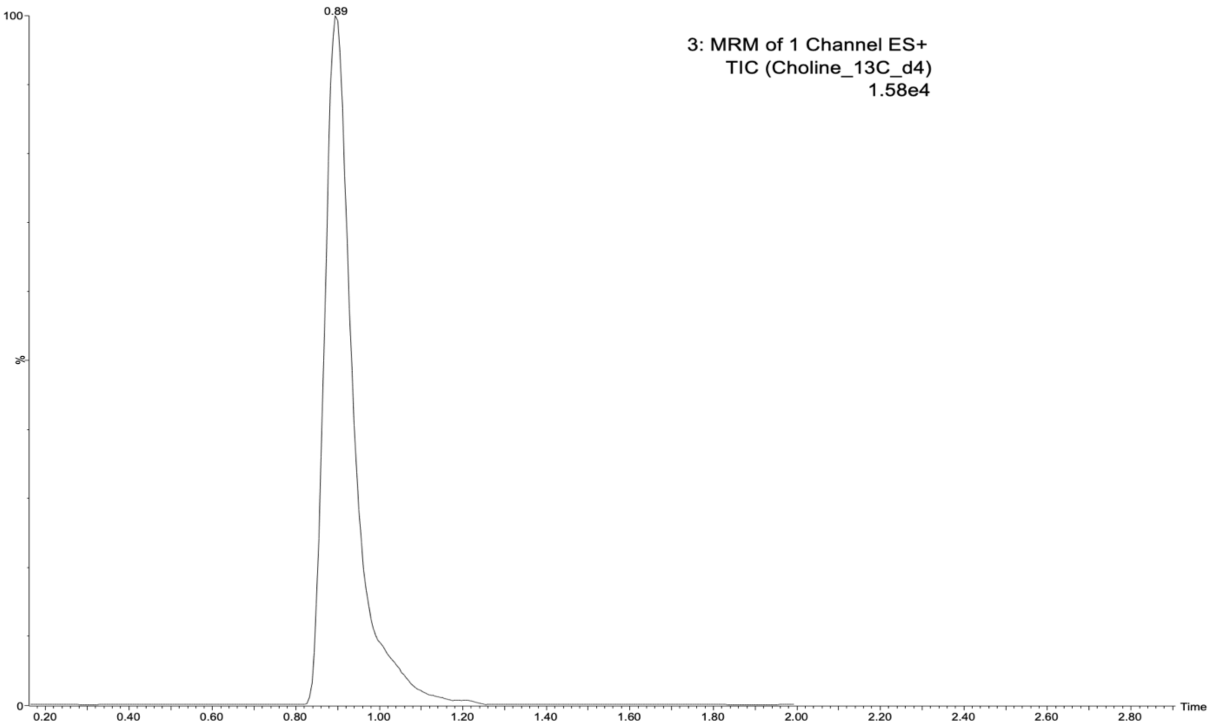


***Supplementary Figure 11****. Representative UPLC-MS/MS chromatogram of internal standard choline-$^{13}$C-d$_4$.*

### Experiment-Specific Methods

**Experiment 2 (section 3.2):** To identify what portion of blueberry tissue exhibits the greatest effect on TMA-lowering activities, whole blueberry, sugar, fiber, and phenolic acid fractions were used in the extended fermentation model (30 hrs as opposed to 24 hrs). Blueberries from two sources (fresh + frozen) were pooled to make a "representative" blueberry sample. Whole blueberries were used to extract phenolic compounds as per methodology described in "*polyphenol extraction*". Digestions were performed as described in the "*general gastrointestinal procedures*". Digestions are scaled from ~1 serving in human volume (1 cup or ~150 g fresh blueberries, ~14.7 g sugar content, ~3.55 g fiber content, and ~1.0125 g phenolic extract content) to an equivalent dose for the *in vitro* digestion final volumes (**Supplementary Table 4**). A "blank" digesta was made with saline in place of food material but contained all other components of the digestion.

**Supplementary Table 4.** Masses of food material for a 50 ml digestion.

| Fraction (*n*=2) | Amount in 1 serving (g) per 2 L upper GI volume | Target mass (mg) in 50 mL digestion | Actual mass (mg) in 50 mL digestion[1] |
|---|---|---|---|
| Whole Blueberry | 150 | 3750 | A: 3.7275, B: 3.7275 |
| Sugar mixture[2] | 14.7 | 367.5 | A: 367.6, B: 367.5 |
| Fiber mixture[3] | 3.55 | 88.75 | A: 88.8, B: 88.7 |
| Phenolic extract | 1.0125 | 25.3125 | A: 25.4, B: 25.9 |

| |
|---|
| [1]letters indicate experimental replicates<br>[2]Comprised of 1.12% glucose, 48.97% glucose, and 49.90% fructose (all wt%)<br>[3]Comprised of 11.27% pectin, 44.36% cellulose, and 44.37% hemicellulose (all wt%) |

Blueberry belongs to the Ericaceae Juss- Health family and the genus *Vaccinium* L.[4]. Blueberries exhibit varying levels of sugar, fiber, and phenolic compounds depending on genotype, growth conditions, post-harvest conditions, and storage environment. Fresh blueberry composition is typically comprised of about 84% water, 9.7% carbohydrate, 0.7% protein, 0.3% fat, and 2.4% fiber[5,6]. A typical serving size of blueberries is about 1 cup (~150 g fresh weight) which contains 21.4 g carbohydrates by difference, 1.09 g of protein, 0.48 g fat, 14.7 g of sugar, and 3.55 g of fiber[6,7]. Of the total sugar content, roughly 0.163 g (1.11%) consist of sucrose, 7.22 g (49.11%) consist of glucose, and 7.36 g (50.06%) consist of fructose[6]. Of the total fiber content, roughly 0.40 g (11.26%) consists of soluble fiber (pectin) and 3.15 g (88.73%) consists of insoluble fiber (cellulose, hemicellulose, lignin)[8].To mimic blueberry's sugar and fiber content, mixtures were made in the same ratios as reported in blueberry and masses corresponding to the total mass of the sugar and fiber fractions were used (see above, footnotes to **Supplementary Table 4).**

Fermentations were carried out as per methodology described in "*general fermentation procedures*". Three different fecal samples were combined with 27 mL of growth media and filtered to remove solids. The 50 mL tube containing the fecal the slurry was left uncapped in the anaerobic chamber for 12 hrs. From 0 to 30 hours, 100 µL of sample was collected at 0, 2, 4, 6, 10, 14, 18, 22, 26, and 30 hr time points, combined with 100 µL of acetonitrile, and frozen at −80°C. To confirm the treatments did not have a cytotoxic or cytostatic effect on fecal bacteria cells, an ATP assay was used at 14 hr and 30 hr time points to monitor cell growth and cell respiration following methodology described in "*BacTiter-Glo Assay*" and "*BacTiter-Glo Analysis".*

Sample extraction and quantification was carried out as described in "*extraction and quantification*". Internal standards for this experiment included choline-1-$^{13}C$-1,1,2,2, -$d_4$ and TMA-$^{13}C_3$-$^{15}N$. Data analysis was carried out as described in "choline and TMA analysis" utilizing a two-way ANOVA test to evaluate the statistical differences between choline-$d_9$ and TMA- $d_9$

**<u>Experiment 3 (section 3.3):</u>** Total phenolic levels were determined for whole blueberry, skin, and pulp fractions using methodology described in "*plant phenolic extraction*" and "*Phenolic analysis: Folin- Ciocalteu assay*". CGA content was determined using a solid phase elution (SPE) and LC/MS quantification as described in "*Solid Phase Elution (SPE)*". Sugar and fiber fractions were included in the digestion and fermentation to compare inhibition levels.

To separate the skin from the pulp, a revised blanching method was used. The whole blueberries were submerged in liquid nitrogen for 10 seconds, removed from the liquid nitrogen and submerged in boiling water for 5 seconds, and then removed from the water and submerged in ice water for 15 seconds. After this process, the peels separated from the pulp with ease. The blueberries were weighed before separation, and both the peel and pulp fractions were weighed. 16.58 g of peel and 107.44 g of pulp were recovered. These values were used to determine masses for the digestion (~13.37% peel and ~86.67% pulp in a serving size of blueberries).

Whole blueberry, skin fraction, and pulp fraction were blended separately using a polytron. The blended fractions were flash frozen and stored at −80°. It is important to note that this method may influence the enzymatic oxidation of polyphenols in the blueberries. When the berries are consumed and chewed, the polyphenols are exposed to polyphenol oxidase (PPO), the enzyme responsible for the oxidation of phenolic compounds. The blanching method used can deactivate PPO, preventing the phenolics from oxidation and leading to effects in the *in vitro* experiment that may not be observed *in vivo*[9].

For the Folin extract, 249.72 mg of whole blueberry, 250.94 mg of peel, and 250.90 mg of pulp were used. To quantify the total phenolics in the extracts, a Folin assay was carried out in a 96 well plate. Each extract underwent a series of dilutions. Starting with the pure extract then diluting 10x, then diluting 100x, etc. A standard curve was made with a 2mg/mL gallic acid mixture with a series of 23- 2/3rds dilutions with a "blank" as pure water. Total phenolics were expressed as mg of gallic acid equivalent (GAE)/ g or whole blueberry (**Table 1**).

To further quantify the relative CGA amounts in the blueberry fractions, the extracts were analyzed using the LC-MS following the SPE (solid phase elution). For the SPE extract, 249.7 mg of whole blueberry, 250.5 mg peel, and 499.6 mg pulp were used. A standard curve was made with a 1mg/mL CGA mixture with a series of dilutions. Samples were plated in triplicate. Total CGA content was expressed as mg CGA/ g whole blueberry (**Table 1**).

Digestions were performed as described in the " *general gastrointestinal procedures*" adopted from Iglesias-Carres *et al.*[1]. Blueberries from two sources (fresh + frozen) were pooled to make a "representative" blueberry sample. Digestions are scaled from ~1 serving in human volume (1 cup or ~150 g fresh blueberries, ~20.05 g peel content, 129.95 g pulp content, ~14.7 g sugar content, and~3.55 g fiber content.) to an equivalent does for the *in* vitro digestion final volumes (**Supplementary Table 5**). A "blank" digesta was made with saline in place of food material but contained all other components of the digestion.

**Supplementary Table 5.** Masses of food material for a 50 ml digestion

| Fraction (*n*=2) | Amount in 1 serving (g) per 2 L upper GI volume | Target mass (mg) in 50 mL digestion | Actual mass (mg) in 50 mL digestion[1] |
|---|---|---|---|
| Whole Blueberry | 150 | 3750 | A: 3770, B: 3750 |
| Peel | 20.05 | 501.25 | A: 501.73, B:501.89 |
| Pulp | 129.95 | 3248 | A: 3260, B:3270 |
| Sugar | 14.7 | 367.5 | A: 366.25, B: 367.24 |
| Fiber | 3.55 | 88.75 | A: 87.44, B: 87.82 |

[1]letters indicate experimental replicates

Fermentations were carried out as per methodology described in "*general fermentation procedures*". Two different fecal samples were combined with 16 mL of growth media and left uncapped in the anaerobic chamber for 12 hrs. The slurry was filtered immediately before the start of the fermentation. The remaining 2 mL of media was used to rinse residual fecal slurry through the filter. The 50 mL tube containing the fecal the slurry was left uncapped in the anaerobic chamber for 12 hrs. From 0 to 24 hours, 100 µL of sample was collected at 0, 2, 4, 6, 8, 10, 12, 16, 20, and 24 hr time points, combined with 100 µL of acetonitrile, and frozen at

−80°C. To confirm the treatments did not have a cytotoxic or cytostatic effect on fecal bacteria cells, an ATP assay was used at 12hr (Figure. 4A.) and 24 hr (Figure. 4B.) time points to monitor cell growth and cell respiration following methodology described in "*BacTiter-Glo Assay*" and "*BacTiter-Glo Analysis".*

Sample extraction and quantification was carried out as described in "*extraction and quantification*". Internal standards for this experiment included choline-1,2-$^{13}C_2$ and TMA-$^{13}C_3$-$d_9$. Data analysis was carried out as described in "*choline and TMA analysis*" displaying kinetic curves for choline- $d_9$ and TMA- $d_9$, utilizing a one-way ANOVA test to evaluate the statistical differences between choline-$d_9$ and TMA- $d_9$ AUCs.

**<u>Experiment 4 (section 3.4)</u>:** Extracts were prepared for each of the 20 different blueberry genotypes as described in "*plant phenolic extraction*". A SPE (solid phase elution) and LC-MS analysis was completed using the extracts as described in "*Solid Phase Elution (SPE) and Analysis*". For the extract, ~250 mg of blueberries from each genotype (*n*=4) were used in four separate replicates. This allowed each replicate to consist of a unique set of blueberries from the same genotype, ensuring a more varied representation of each genotype. (**Supplementary Table 6**). A standard curve was made with a 4mg/mL CGA mixture with a series of 35- 2/3rds dilutions with the 35$^{th}$ well as pure water. Samples were plated in *n*=4. Total CGA content was expressed as mg CGA/ 100g whole blueberry (**Table 2**.). CGA content is used to determine four of the highest CGA content and four of the lowest CGA content blueberry genotypes to use in the fermentation model.

**Supplementary Table 6.** Masses for genotype extracts

| **Gentotype (*n*=4)** | **Mass (mg)[1]** |
|---|---|
| DxJ 197 | A: 252, B: 252, C: 251.3, D: 251.8 |
| DxJ 148 | A: 251, B: 251.5, C: 253, D: 253.1, |
| DxJ 112 | A: 247.3, B: 253.7, C: 248.9, D: 251.6 |
| DxJ 041 | A: 248.8, B: 250.7, C: 249.5, D: 251.8 |
| DxJ 125 | A: 249.1, B: 251.6, C: 253.2, D: 249 |
| DxJ 124 | A: 251.1, B: 249.9, C: 252, D: 248.6 |
| DxJ 023 | A: 252.9, B: 251.7, C: 251.6, D: 249.4 |
| DxJ 102 | A: 251.1, B: 251.7, C: 250, D: 251 |
| DxJ 038 | A: 250.6, B: 252.5, C: 250.6, D: 252.8 |
| DxJ 149 | A: 253, B: 249.7, C: 252.7, D: 252.1 |
| DxJ 002 | A: 252, B: 250.2, C: 251.3, D: 249.5 |
| DxJ 033 | A: 250.5, B: 249.9, C: 251.6, D: 251.1 |
| PI 554857 | A: 251.3, B: 253, C: 250.9, D: 250.5 |
| PI 554856 | A: 251.5, B: 250.6, C: 250.5, D: 251.1 |
| PI 554764 | A: 253.5, B: 250.8, C: 251.5, D: 250.7 |
| PI 554846 | A: 250.6, B: 252.9, C: 250.1, D: 251 |
| PI 554849 | A: 253, B: 251.3, C: 251.1, D: 252.9 |
| PI 638335 | A: 249.9, B: 250.3, C: 253.5, D: 253.6 |
| PI 267851 | A: 249.9, B: 248.8, C: 252.7, D: 249.8 |
| PI 296399 | A: 252.3, B: 252.3, C: 251.4, D: 251.6 |

[1]letters indicate experimental replicates

The highest four CGA content genotypes (DxJ 033, PI 267851, DxJ 002, PI 296399) and the lowest four CGA content genotypes (PI 554856, PI 554857, DxJ 197, DxJ 148) were peeled by hand, blended with a polytron (VWR 200), and weighed for a 15 mL digestion. Digestions were performed as described in the "general gastrointestinal procedures". Digestions are scaled from ~1 serving in human volume (150 g FW providing ~20.05 g peel content) to an equivalent dose for the *in vitro* digestion final volumes (**Supplementary Table 7).** A "blank" digesta was made with saline in place of food material but contained all other components of the digestion.

**Supplementary Table 7.** Masses of peel material used in 15 ml digestion

| Genotype (*n*=2) | CGA level | Amount in 1 serving (g) per 2 L upper GI volume | Target mass (mg) in 50 mL digestion | Actual mass (mg) in 50 mL digestion |
|---|---|---|---|---|
| DxJ 197 | Low | 20.05 | 150.375 | A: 152.4, B: 149.1 |
| DxJ 148 | Low | | | A: 150.5, B: 149.8 |
| DxJ 002 | High | | | A: 150.6, B: 150.0 |
| DxJ 033 | High | | | A: 149.6, B: 151 |
| PI 554857 | Low | | | A: 154.3, B: 151.8 |
| PI 554856 | Low | | | A:151.7, B: 149.7 |
| PI 267851 | High | | | A: 151.9, B: 151.6 |
| PI 296399 | High | | | A: 151.8, B: 150.6 |

Fermentations were carried out as per methodology described in "general fermentation procedures". Three different fecal samples were combined with 27 mL of growth media and left uncapped in the anaerobic chamber for 12 hrs. The slurry was filtered immediately before the start of the fermentation. The remaining 2 mL of media was used to rinse residual fecal slurry through the filter. The 50 mL tube containing the fecal the slurry was left uncapped in the anaerobic chamber for 12 hrs. From 0 to 24 hours, 100 µL of sample was collected at 0, 2, 4, 6, 8, 10, 12, 16, 20, 24 hr time points, combined with 100 µL of acetonitrile, and frozen at −80°C. To confirm the treatments did not have a cytotoxic or cytostatic effect on fecal bacteria cells, an ATP assay was used at 12hr and 24 hr time points to monitor cell growth and cell respiration following methodology described in "BacTiter-Glo Assay" and "BacTiter-Glo Analysis".

Sample extraction and quantification was carried out as described in "*extraction and quantification*". Internal standards for this experiment included choline-1,2-$^{13}C_2$ and TMA-$^{13}C_3$-$d_9$. Data analysis was carried out as described in "choline and TMA analysis" displaying kinetic curves for choline- $d_9$ and TMA- $d_9$. Utilizing a one-way ANOVA test to evaluate the statistical differences between choline-$d_9$ and TMA- $d_9$ AUCs.

Fruit size determination. Average genotype size was measured to determine whether CGA content correlates with size of the blueberry. If smaller blueberries contain a higher amount of CGA, a smaller quantity will need to be consumed to achieve the health benefits associated with blueberries. 15 blueberries from each genotype were selected at random and the size (diameter) was measured in mm with an electric Mitutoyo digimatic caliber and recorded.

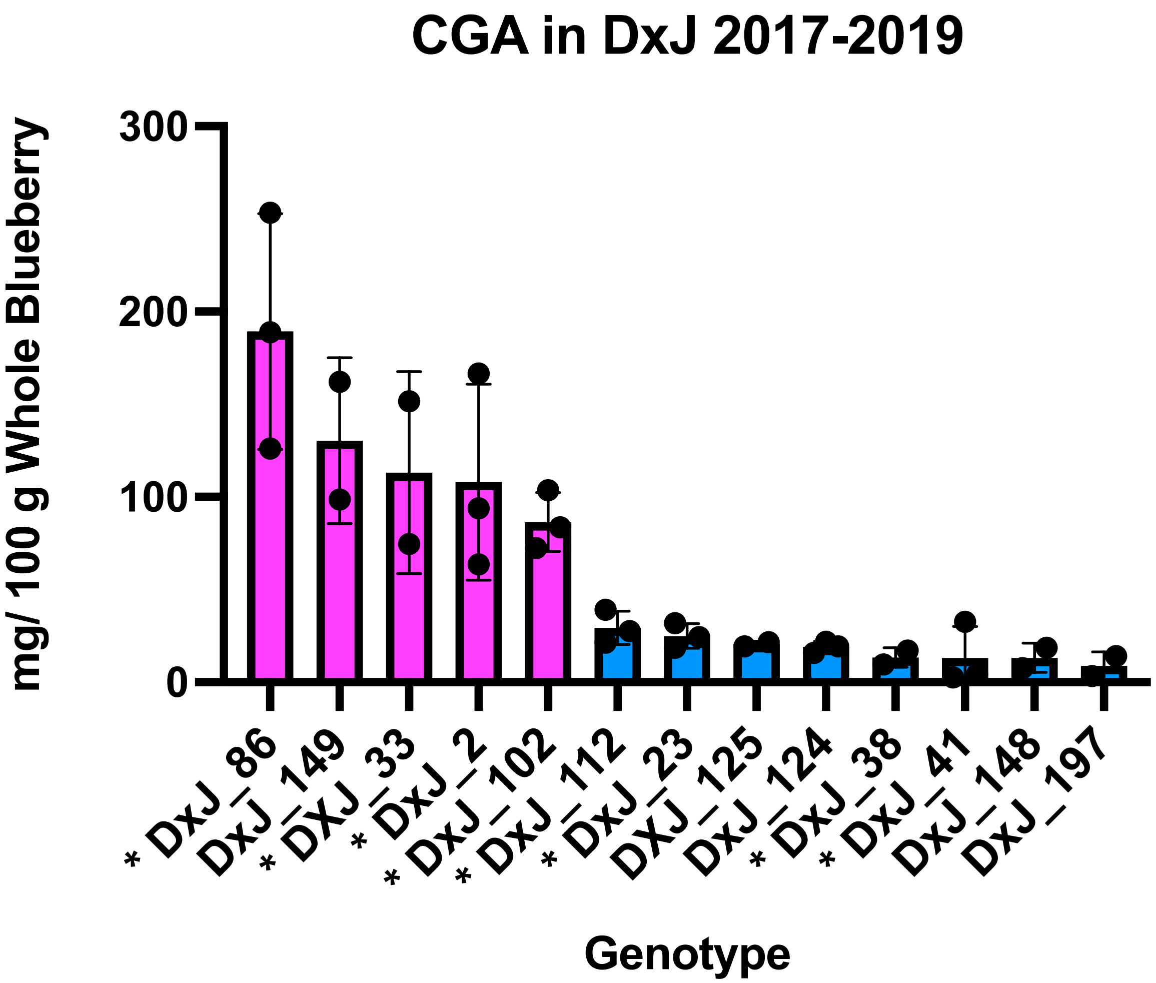


**Supplementary Figure 12.** Chlorogenic acid (CGA) content on a fresh weight (FW) basis in blueberries from a Draper x Jewel (DxJ) biparental mapping population (2017-2019). Black dots represent mean of individual years with error bars representing standard error of the mean (SEM) across years. Genotypes with asterisks were selected for initial digestion and fermentation experiment. Data were adapted from Mengist *et al.* [10]with permission (https://creativecommons.org/licenses/by/4.0/) .

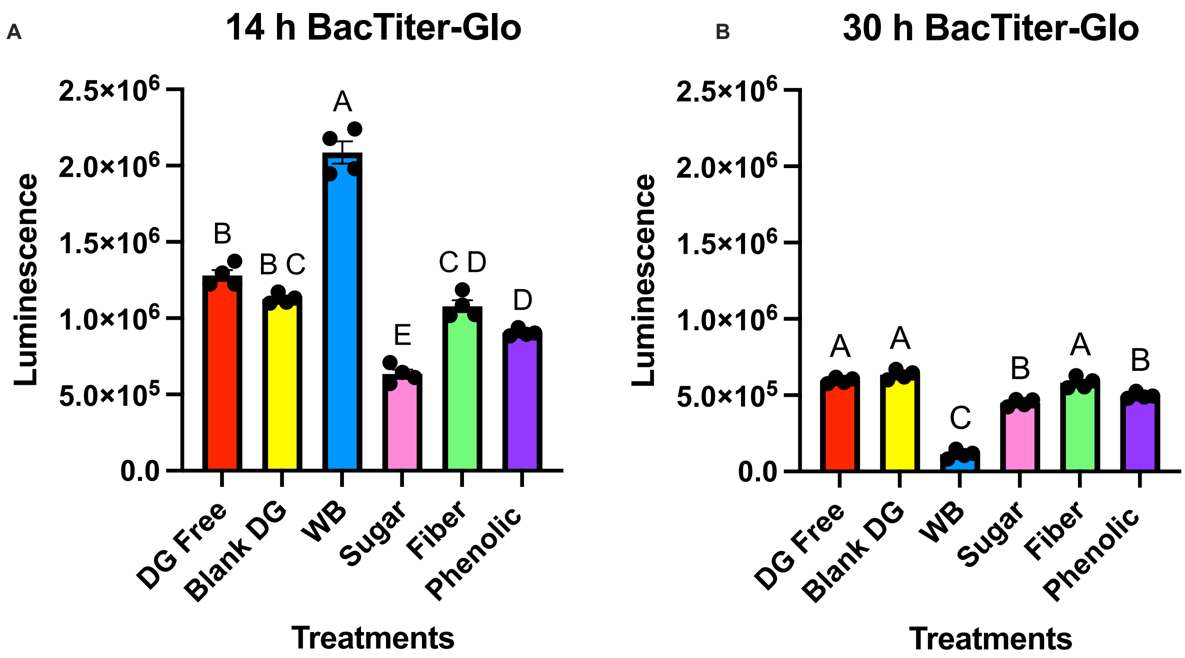


**Supplementary Figure 13.** BacTiter-Glo luminescence reading at 14 hr (A) and 30 hr (B) for e*xperiment 2: Determination of the impacts of phenolic (CGA) and non-phenolic components (sugar, fiber, skin, pulp) of blueberries on TMA production. Treatments:* Digesta free (DG free), Blank DG (blank digesta), Whole blueberry (WB), Sugar, Fiber, and Phenolic. Using one-way ANOVA and Tukey's multiple comparison to determine statistical differences between treatments. Different letters notate a statistical difference (P<0.05). Data represent mean ± SEM from *n*=4.

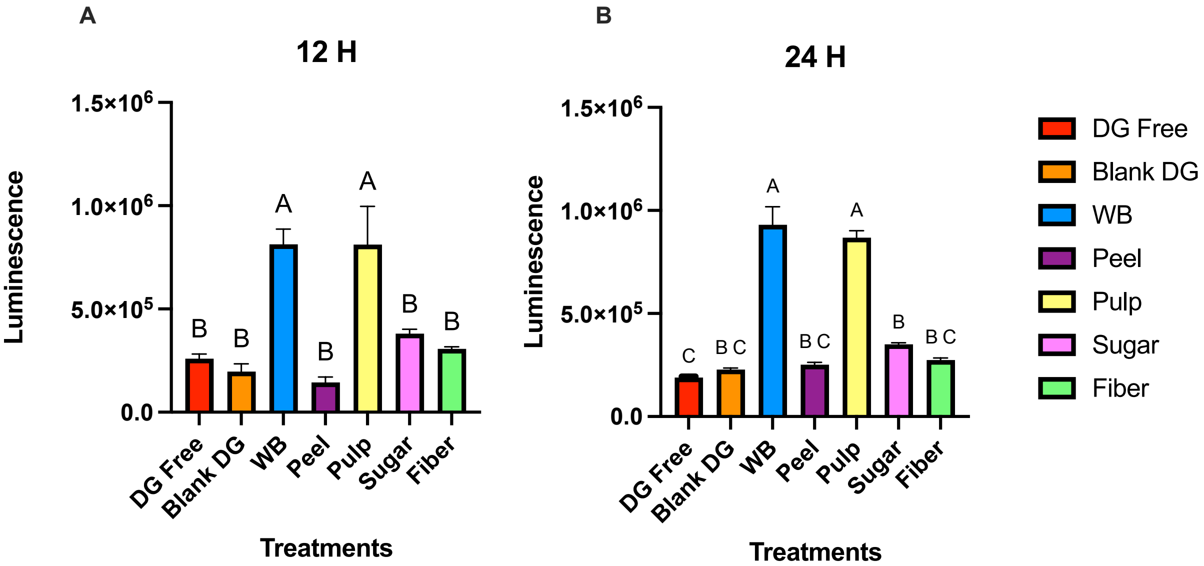


**Supplementary Figure 14**. BacTiter-Glo luminescence reading at 12 hr (A) and 24 hr (B) for *experiment 3. Identification of blueberry components with the greatest TMA inhibition potential:* Treatments: Digesta (DG), whole blueberry (WB): DG free, blank DG, WB, Peel, Pulp, Sugar, and Fiber. Using one-way ANOVA and Tukey's multiple comparison to determine statistical differences between treatments. Different letters notate a statistical difference ($P<0.05$). Data represent mean ± SEM from $n$=6.

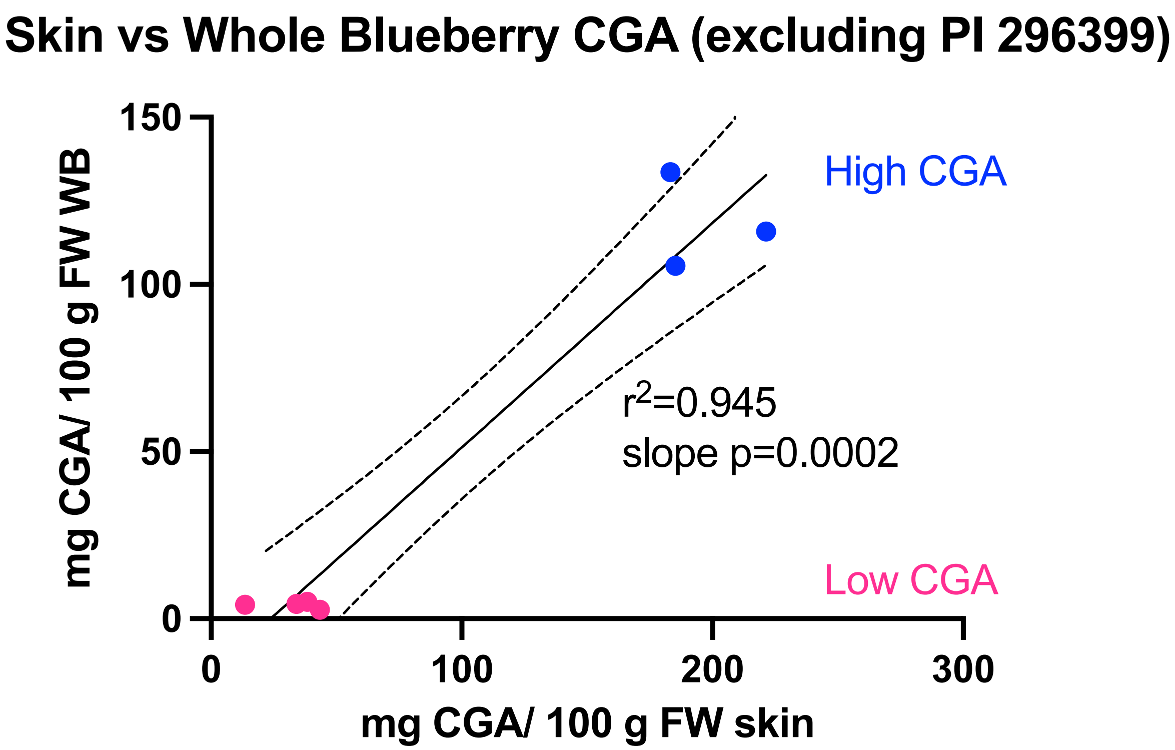


**Supplementary Figure 15.** Correlation of chlorogenic acid (CGA) content in skins and whole fruits in the selected genotypes with extreme CGA content, excluding PI 296399. Dots represent means.

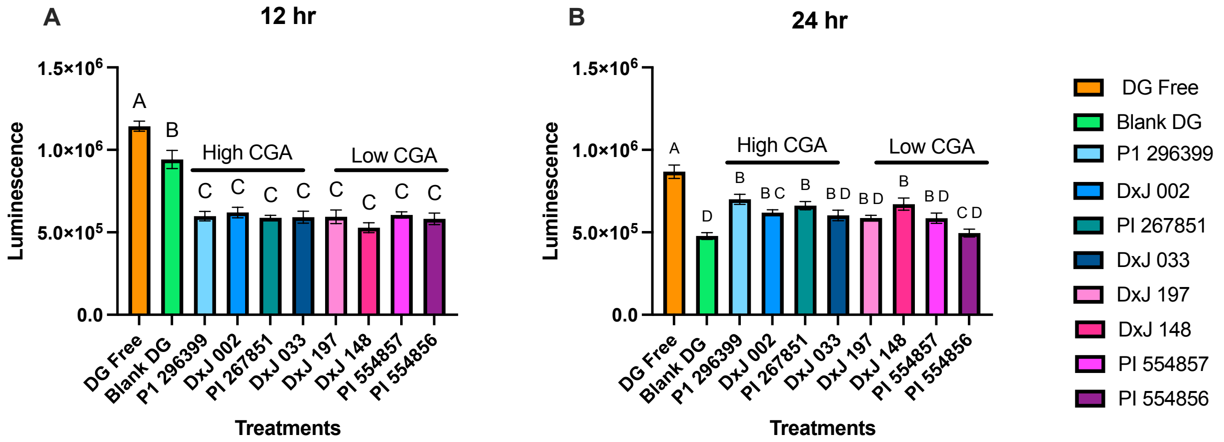


**Supplementary Figure 16.** BacTiter-Glo luminescence of choline-$d_9$ concentration (A) and TMA-$d_9$ (B) production in fermentation (0-24 hr) for *experiment 4. Determining whether CGA content correlates with reduced TMA production in multiple blueberry accessions from a genetic diversity population with a large spread of CGA in our ex vivo-in vitro human fecal fermentation model.* Treatments: digesta (DG): DG free, Blank DG, PI 296399 (high CGA), DxJ 002 (High CGA), PI 267851 (High CGA), DxJ 033 (High CGA), DxJ 197 (Low CGA), DxJ 148 (Low CGA), PI 554857 (Low CGA), PI 554856 (Low CGA). using one-way ANOVA and Tukey's multiple comparison to test for treatment effects: Data represent mean ± SEM from $n$=6.

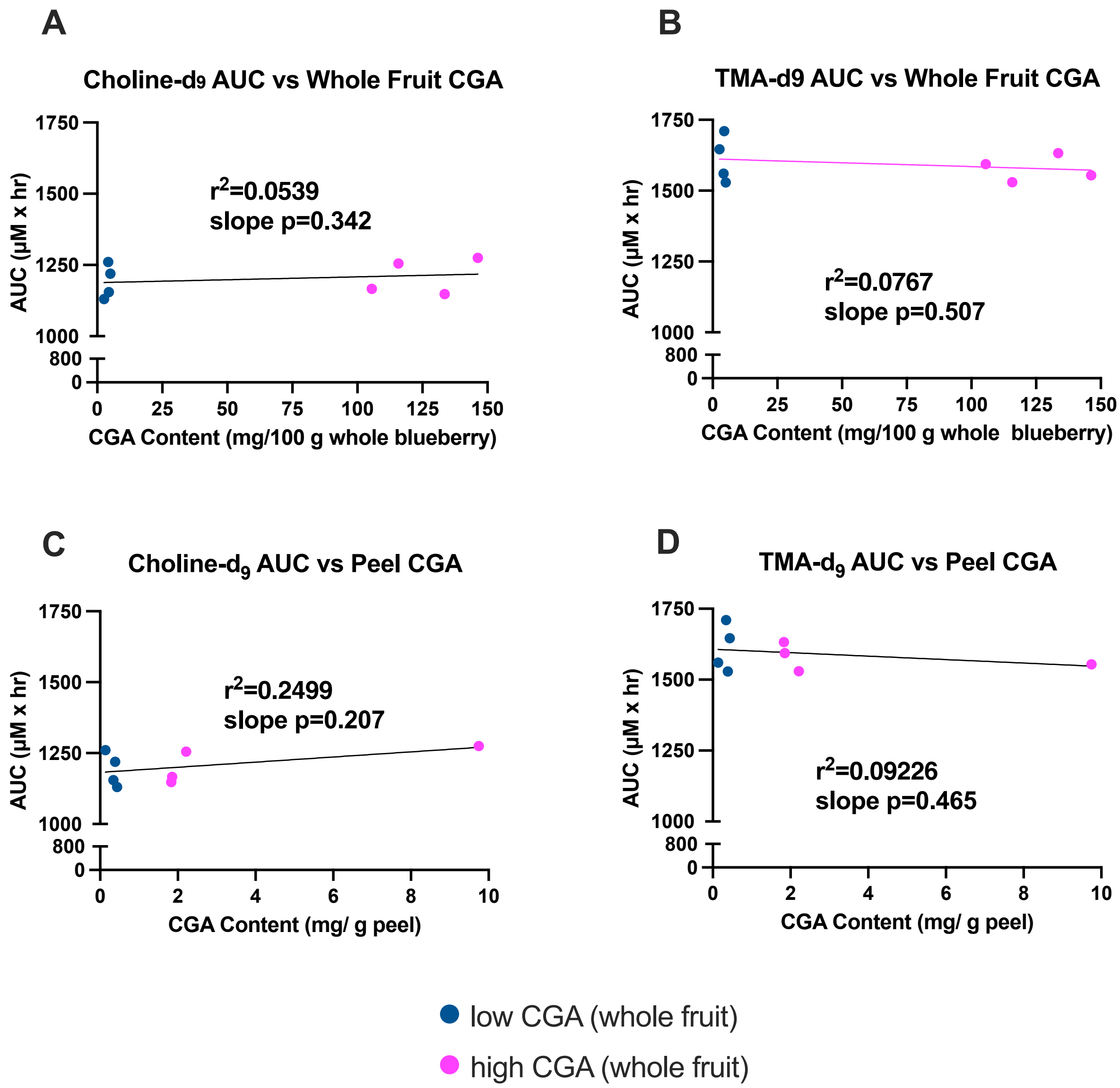


**Supplementary Figure 17.** Correlations between chlorogenic acid (CGA) content in whole blueberry (A-B) or peel (C-D) with fermentation area-under-the-curve (AUC) values for choline-$d_9$ (A, C) or TMA-$d_9$ (B, D). Points represent mean values for each individual genotype.

### REFERENCES

1 L. Iglesias-Carres, L. A. Essenmacher, K. C. Racine and A. P. Neilson, *Nutrients*, 2021, **13**, 1466.

2 Development of a genetic framework to improve the efficiency of bioactive delivery from blueberry - PubMed, https://pubmed.ncbi.nlm.nih.gov/33057109/, (accessed January 1, 2025).

3 In vitro evidences of the globe artichoke antioxidant, cardioprotective and neuroprotective effects - ScienceDirect,

https://www.sciencedirect.com/science/article/pii/S1756464623002748?via%3Dihub, (accessed January 10, 2024).
4 USDA Plants Database, https://plants.usda.gov/home/classification/65446, (accessed February 24, 2024).
5 A. Michalska and G. Łysiak, *Int. J. Mol. Sci.*, 2015, **16**, 18642–18663.
6 FoodData Central, https://fdc.nal.usda.gov/fdc-app.html#/food-details/171711/nutrients, (accessed February 24, 2024).
7 Blueberries, raw, 1 cup - Health Encyclopedia - University of Rochester Medical Center, https://www.urmc.rochester.edu/encyclopedia/content.aspx?contenttypeid=76&contentid=09050-1, (accessed March 26, 2024).
8 A. Rashidinejad, in *Nutritional Composition and Antioxidant Properties of Fruits and Vegetables*, ed. A. K. Jaiswal, Academic Press, 2020, pp. 467–482.
9 R. Moscetti, F. Raponi, D. Monarca, G. Bedini, S. Ferri and R. Massantini, *Int. J. Food Sci. Technol.*, 2019, **54**, 403–411.
10 M. F. Mengist, M. H. Grace, J. Xiong, C. D. Kay, N. Bassil, K. Hummer, M. G. Ferruzzi, M. A. Lila and M. Iorizzo, *Front. Plant Sci.*
